# Tumor-infiltrating nerves create an electro-physiologically active microenvironment and contribute to treatment resistance

**DOI:** 10.1101/2020.04.24.058594

**Authors:** Attila Kovacs, Daniel W. Vermeer, Marianna Madeo, Hunter D. Reavis, Samuel J. Vermeer, Caitlin S. Williamson, Alex Rickel, Jillian Stamp, Christopher T. Lucido, Jacob Cain, Maria Bell, Mark Morgan, Ju-Yoon Yoon, Marilyn A. Mitchell, Natalia Tulina, Sarah Stuckelberger, Anna Budina, Dalia K. Omran, Euihye Jung, Lauren E. Schwartz, Tuany Eichwald, Zhongkui Hong, Jill Weimer, Jody E. Hooper, Andrew K. Godwin, Sebastien Talbot, Ronny Drapkin, Paola D. Vermeer

**Affiliations:** Sanford Research, 2301 East 60^th^ St North, Sioux Falls, SD 57104, USA; Penn Ovarian Cancer Research Center, Department of Obstetrics and Gynecology, Division of Gynecologic Oncology, University of Pennsylvania, Perelman School of Medicine, 421 Curie Blvd, Philadelphia, PA 19104, USA; Biomedical Engineering Program, University of South Dakota, 4800 North Career Ave, Sioux Falls, SD 57107, USA; Sanford Gynecologic Oncology, Sanford Health, 1309 West 17^th^ St, Sioux Falls, SD 57104, USA; Department of Pathology and Laboratory Medicine, University of Pennsylvania, Perelman School of Medicine, 3400 Spruce St, Philadelphia, PA 19104, USA; Department of Pathology and Laboratory Medicine, University of Kansas Medical Center, 3901 Rainbow Blvd, Kansas City, KS 66160, USA; Department of Pathology, Johns Hopkins University School of Medicine, 600 North Wolfe Street, Baltimore, MD 21287, USA; Department of Pharmacology and Physiology, University of Montreal, 2900 Edouard-Montpetit, Montreal (Quebec) H3T 1J4

**Keywords:** innervation, ovarian cancer, chemotherapy, extracellular vesicles, TRPV1, sensory, micro-electrode array

## Abstract

Patients with densely innervated tumors do poorly as compared to those with sparsely innervated disease. Why some tumors heavily recruit nerves while others do not, remains unknown as does the functional contribution of tumor-infiltrating nerves to cancer. Moreover, while patients receive chemotherapeutic treatment, whether these drugs affect nerve recruitment has not been tested. Using a murine model of ovarian cancer, we show that tumor-infiltrating sensory nerves potentiate tumor growth, decrease survival, and contribute to treatment resistance. Furthermore, matched patient samples show significantly increased tumor innervation following chemotherapy. *In vitro* analysis of tumor-released extracellular vesicles (sEVs) shows they harbor neurite outgrowth activity. These data suggest that chemotherapy may alter sEV cargo, endowing it with robust nerve recruiting capacity.

## INTRODUCTION

A growing body of evidence supports the importance of tumor innervation in cancer [1, 2]. For instance, genetic, chemical and surgical ablations of tumor-infiltrating nerves in cancer models demonstrate active roles for nerves in disease initiation and progression [3-5]. Evidence for the recruitment of central nervous system neural progenitors to tumors in mice further emphasizes the existence of intricate interactions between tumors and the nervous system [6]. In addition, neurotrophic factors and axonal guidance molecules are pro-tumorigenic [7-11] while neurotransmitter receptor blockade is anti-tumorigenic [12-17]. Together, these data suggest the nervous system is not a bystander but an active participant in cancer and indicate that extensive tumor innervation contributes to aggressive disease [1].

While communication between cancer and the nervous system is appreciated, the potential that extracellular vesicles (EVs, vesicles released by cells) are vehicles of this communication was only recently discovered. Three reports in squamous cell carcinomas show that tumor-released small EVs (sEVs) promote innervation in cancer [18-20]. Moreover, highly innervated tumors grow faster and are more metastatic than sparsely innervated disease and sEVs directly contribute to this phenotype [18-20]. Based on these findings, we assessed innervation in other solid tumors (breast, prostate, pancreatic, lung, liver, ovarian and colon) and found that all are innervated. Two recent studies focused our efforts on ovarian cancer. The first shows that loss of monoubiquitinated histone H2B (H2Bub1), an important early event in the evolution and progression of high-grade serous ovarian carcinoma (HGSOC), alters chromatin accessibility thereby activating signaling pathways that contribute to disease progression (23); a dominant signature emerged, that of axonal guidance/neurotrophin/synaptic signaling. This signature, together with an *in silico* analysis of gene expression data by Yang *et al* (24), and our finding of tumor-infiltrating nerves in HGSOCs, prompted us to better define innervation in this lethal disease.

Epithelial ovarian cancer, the fifth most common cancer in women, is the most lethal gynecologic malignancy [21]. Worldwide, nearly 300,000 women are diagnosed with this disease annually; 150,000 will succumb within the first year [22](Wild CP, Stewart B, Weiderpass E and Stewart BW (2020) *World Cancer Report* Lyon: IARC Press). Ovarian cancer is a heterogeneous disease with multiple histologic subtypes. HGSOC accounts for over 70% of cases and the majority of deaths. Surgical de-bulking followed by platinum-based chemotherapy remains standard treatment for HGSOC. While initially effective, the majority of patients progress and succumb to recurrent, chemotherapy-resistant disease [21]. These dismal statistics reflect a poor mechanistic understanding of progression in ovarian cancer. Here we use orthogonal approaches, including mouse models and human tissues, to show that HGSOCs are innervated by sensory nerves that remain functional at the tumor bed. Tumor-released sEVs mediate nerve recruitment and depletion of these nerves leads to decreased tumor growth with improved response to chemotherapy. Importantly, we show that malignant tumors exhibit measurable electrical activity that can be pharmacologically attenuated. Finally, we provide evidence that chemotherapy exacerbates tumor innervation and contributes to aggressive tumor biology. Together, our data show that tumors are innervated by an sEV-mediate process that contributes to disease progression and response to therapy. The ability to impede this process may represent a novel therapeutic opportunity for ovarian cancer and other solid tumors.

## RESULTS

### Sensory nerve twigs innervate tumors

The presence of nerves within HNSCC and cervical cancer patient samples were recently identified by IHC staining for the pan-neuronal marker, β-III tubulin. These tumor-infiltrating nerves are IHC positive for the transient receptor potential vanilloid type 1 channel (TRPV1), a nociceptive sensory marker, but negative for tyrosine hydroxylase (TH, sympathetic marker) and vasoactive intestinal polypeptide (VIP, parasympathetic marker) [18, 19]. To define if other solid tumors are similarly innervated, we surveyed a collection of cancers in a similar fashion. Ten samples/tumor type were scored for innervation by four independent scorers. Similar to HNSCC and cervical cancers, breast, prostate, pancreatic, lung, liver, ovarian and colon cancers harbor β-III tubulin positive nerve fibers (Figure 1A-G). While scoring of tumor-infiltrating nerves was variable, all tumor types analyzed were innervated (Figure 1H). The recent description of a neuronal signature in HGSOC [23, 24] prompted us to focus on defining innervation in this tumor type.

**Figure 1.**
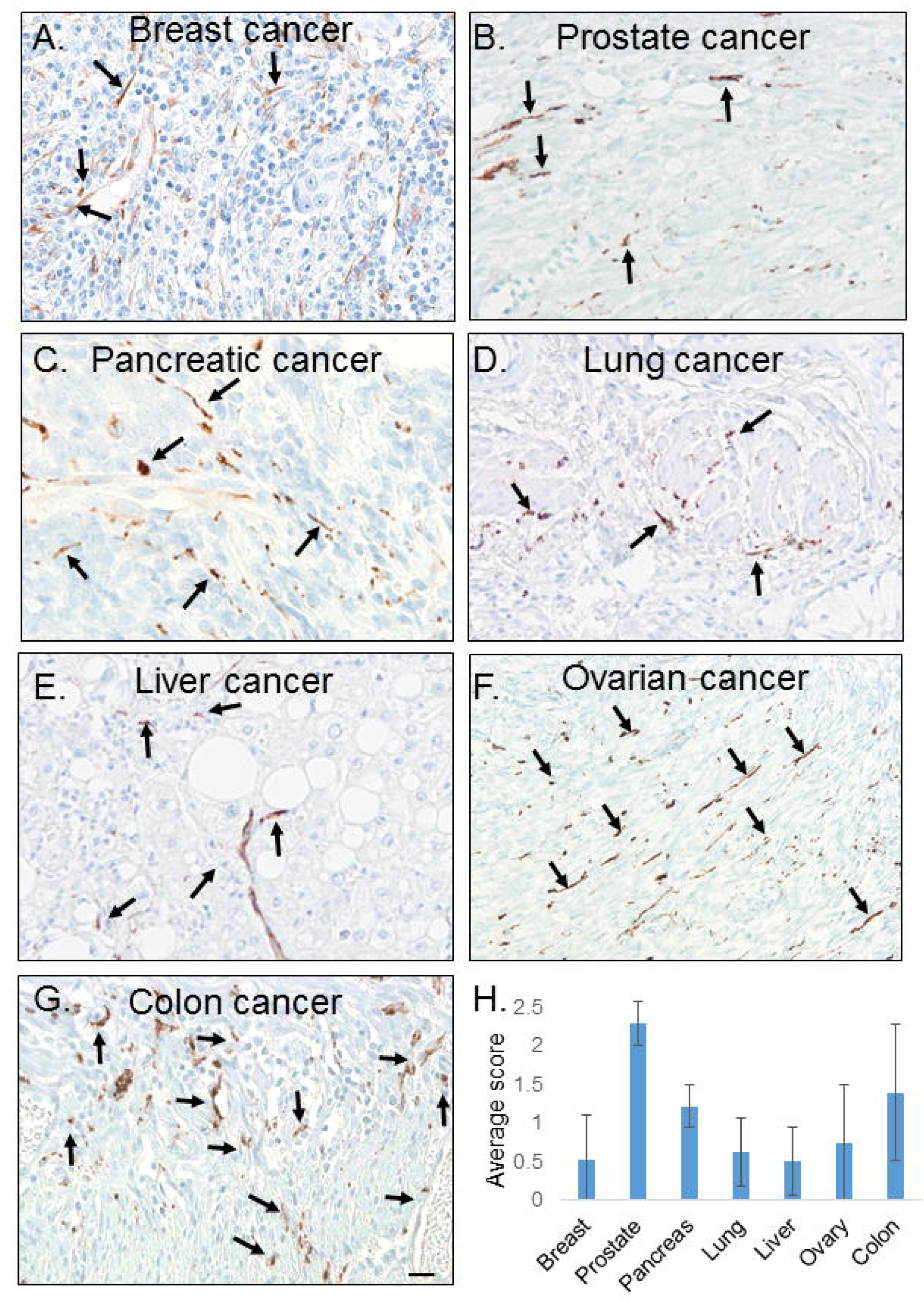
Innervation in solid tumors. A-G) Bright field images of indicated tumors IHC stained for β-III tubulin (brown, arrows; n=10 tumors/type except ovarian with n=30). Light blue, counterstain. Scale bar, 20µm. H) Average innervation score/tumor type. All patient samples were scored for nerve twigs by four independent evaluators, each scored 5 random 20X magnification images/sample. Scoring averages are graphed; standard deviation as error bars.

Recent molecular studies demonstrate that HGSOCs are derived from fallopian tube secretory cells [25-29]; thus, normal tissue controls included normal fallopian tubes and ovaries. IHC staining shows that normal fallopian tube contains TH (sympathetic) positive nerve bundles (Figure 2A, open arrowheads) that are negative for TRPV1 (sensory) and VIP (parasympathetic); scant single nerve fibers (Figure 2A, β-III tubulin positive, small filled arrowheads) are also evident. Normal ovary is similarly innervated with TH positive, TRPV1 and VIP negative nerve bundles (Figure 2A, open arrowheads). While not all HGSOC cases harbored the same extent of nerves, the staining in those that did was in contrast to that of normal tissues; these nerves were TRPV1 positive but negative for TH and VIP (Figure 2A, arrows). Positive controls for VIP, TRPV1 and TH IHC can be found in Supplemental Figure 1A-C. To further validate the presence of tumor-infiltrating nerves, we immunofluorescently stained patient samples for neurofilament, another neuronal marker (Supplemental 1D). To confirm that β-III tubulin positive twigs were TRPV1 positive, tumors were double immuno-stained to demonstrate their co-localization (Figure 2B). Since the type of innervation (sensory) in HGSOC differs from that in normal fallopian tube and ovary (sympathetic), these data suggest that HGSOCs obtain sensory nerves as a consequence of disease rather than by default.

**Figure 2.**
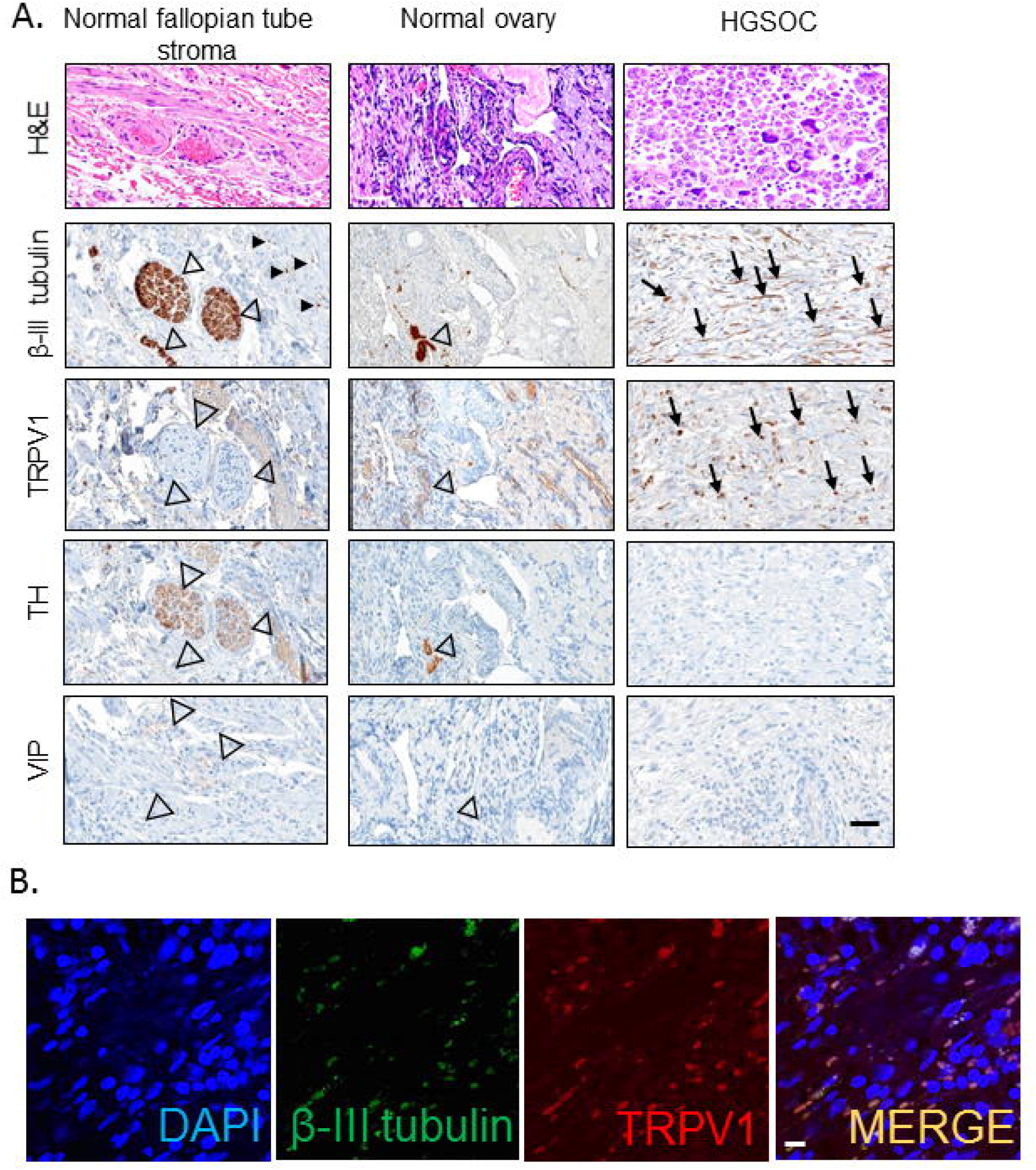
HGSOC innervation. A) Representative bright field images of normal fallopian tube (n=10), normal ovary (n=10) and HGSOC samples (n=30) histochemically stained with hematoxylin and eosin or immunohistochemically stained as indicated. Large arrowheads, nerve bundles; small black arrowheads and small arrows, nerve twigs; scale bar, 10µm. B) Representative *en face* confocal images of HGSOC sample double immunofluorescently stained as indicated. N=8 patient samples stained. Scale bar, 10µm. Brightness was increased on all images in all lasers; these changes were made to the entire image.

We noted that many HGSOC tumor cells themselves were also positive for β-III tubulin; some samples exhibiting robust immunostaining (Supplemental Figure 2A), others with variable staining (Supplemental Figure 2B) and still others predominantly negative for β-III tubulin (Supplemental Figure 2C). While the significance of this staining remains unclear, correlations with aggressive disease and poor survival exist [30, 31]. Given our interest in tumor innervation, however, we focused only on β-III tubulin positive nerves. We also noted the proximity of nerves to tumor cells; in some samples, nerves were embedded within islands of tumor cells (Supplemental Figure 2D, E) while in others, they were present in the stroma, in close proximity to tumor (Supplemental Figure 2F, G).

### Cell-derived sEVs harbor neurite outgrowth activity

Tumor-released sEVs have previously been shown to harbor neurite outgrowth activity that promotes tumor innervation [18, 19]. Compromising tumor sEV release, genetically or pharmacologically, attenuates tumor innervation *in vivo*, emphasizing an active role of sEVs in this process [19]. The tumor-infiltrating nerves evident in HGSOCs are similar to those previously identified. Therefore, we tested whether sEVs from HGSOC cell lines possess neurite outgrowth activity. sEVs from conditioned media were purified by differential ultracentrifugation as previously described [19]. To validate our methodology, we analyzed purified sEVs by atomic force microscopy and found their size (84-130nm) was consistent with sEVs (Figure 3A). As a further validation, purified sEVs from various HGSOC and control fallopian tube cell lines were analyzed by western blot for sEV markers, CD9 and CD81 (Figure 3B) [32]. Moreover, since some published work support a more stringent isolation of sEVs, we further purified some preparations by Optiprep density gradient centrifugation [33, 34]. Consistent with previous findings, CD9 and CD81 positive sEVs were present in fraction 8 (Supplemental Figure 3A) as well as in sEVs purified by differential ultracentrifugation alone (“crude”) [19]. Satisfied that HGSOC cell lines release sEVs and our methodology successfully purifies them from conditioned media, we tested their axonogenic potential utilizing PC12 cells, a rat pheochromocytoma cell line, as a surrogate for neurite outgrowth activity. When appropriately stimulated (e.g. nerve growth factor, NGF), PC12 cells differentiate into neuron-like cells and extend neurites [19, 35, 36]. Given that HGSOCs arise from fallopian tube secretory cells [25, 37, 38], we purified sEVs from the conditioned media of an isogenic set of cell lines as follows. The FT33-Tag cell line was developed from normal human fallopian tube secretory cells by stable expression of large T-antigen (FT33-Tag) [39]. Two transformed cell lines were derived from FT33-Tag that stably express either Myc (FT33-Myc) or Ras oncogenes (FT33-Ras). Importantly, when implanted in NSG (immune incompetent) female mice, FT33-Tag cells are not tumorigenic while FT33-Myc and FT33-Ras are [39]. sEVs were purified from the three isogenic FT33 cell lines, analyzed by nanosight particle analysis (Supplemental Figure 3B) and quantified. Equal amounts of sEVs as measured by protein assay were used to stimulate PC12 cells. Forty-eight hours later, PC12 cells were immunostained for β-III tubulin (Figure 3C) and quantitative microscopy determined the extent of fluorescent β-III tubulin stained neurites/well as a measure of sEV-mediated neurite outgrowth. NGF treatment drives a robust response and experimental values were normalized to this control. We found that FT33-Tag sEVs were unable to induce neurite outgrowth of PC12 cells above the negative control while those from FT33-Myc and FT33-Ras cells induced robust neurite outgrowth (Figure 3C, D). To verify that sEV-mediated neurite outgrowth activity is not unique to this isogenic series of cell lines, sEVs from several routinely used HGSOC cell lines (KURAMOCHI, OVCAR-3, OVCAR-4, FU-OV-1 and OV-90) and three additional control FT lines (FT190, FT194, FT246) [25, 39, 40] were similarly purified and tested. Treatment of PC12 cells with HGSOC sEVs induced significant neurite outgrowth as compared to sEVs from control FT lines (Supplemental Figure 3C, D). These data suggest that sEVs released by HGSOC cell lines possess neurite outgrowth activity which is absent from normal fallopian tube epithelial sEVs.

**Figure 3.**
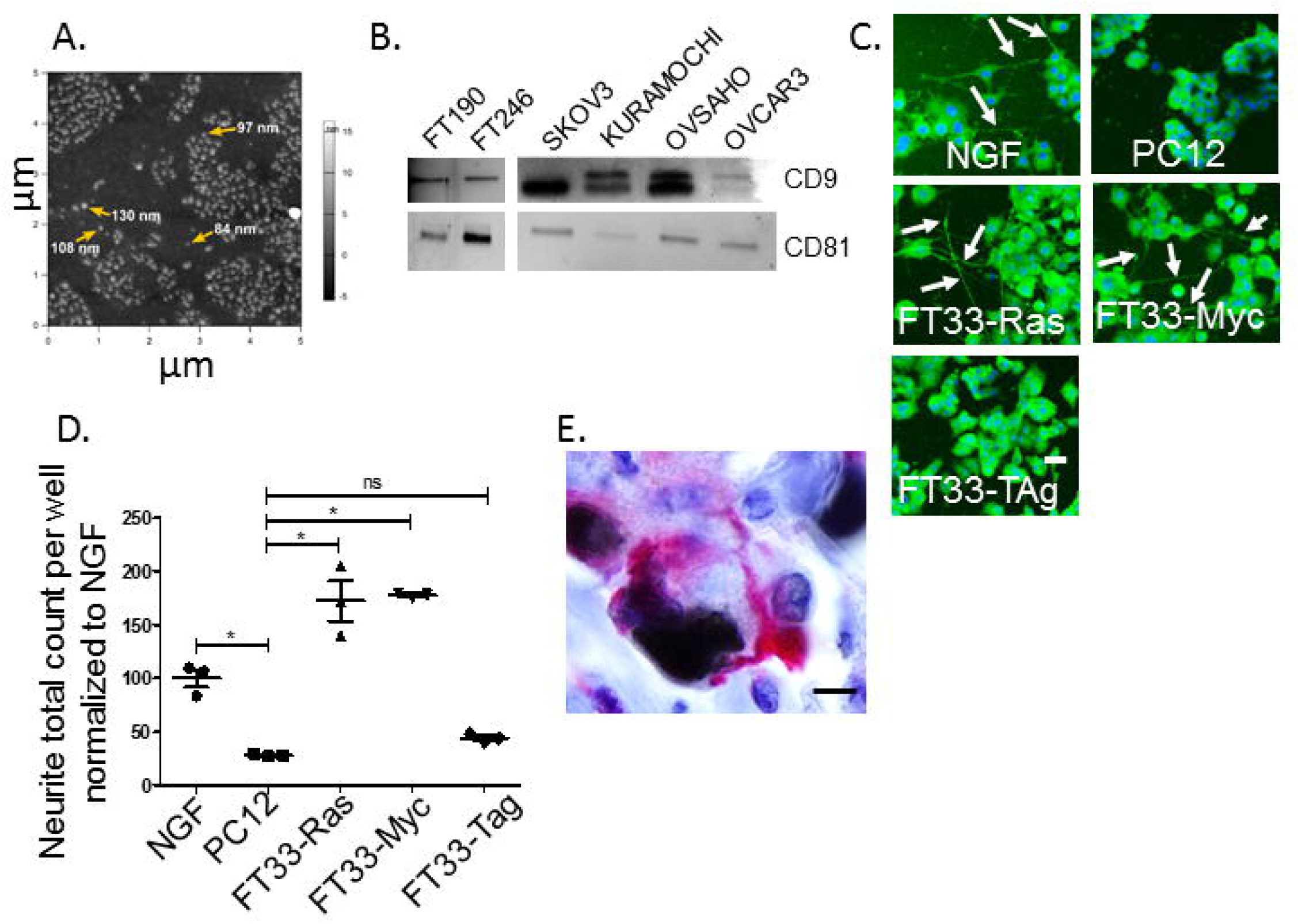
HGSOC sEVs harbor neurite outgrowth activity. A) Topographic image of HGSOC sEVs analyzed by atomic force microscopy. Yellow arrows indicate sized sEVs. B) Western blot analysis of sEVs from the indicated cell lines for CD9 and CD81. C) Representative *en face* fluorescent images of β-III tubulin stained (green) PC12 cells following stimulation with sEVs from the indicated cell lines or with recombinant NGF. Unstimulated PC12 cells (PC12), negative control. Scale bar, 10 µm. D) Quantification of total β-III tubulin positive neurites per well for PC12 cells stimulated with sEVs from the indicated cell lines. PC12 cells stimulated with 50ng/ml NGF (positive control); unstimulated PC12 cells (negative control). N=4 wells/condition (technical replicates). The experiment was repeated at least two times (biological replicates). One way ANOVA with post-hoc Fisher’s Least Significant Difference (LSD) test was used for statistical analysis. LSD p values reported. Error bars indicate standard deviation. Center value used was the mean. *, p<0.05; ns, not significant. The variance between groups compared is similar. E) Representative bright field image of double IHC stained human ovarian tumor for Pax8 (brown) and β-III tubulin (pink), scale bar, 20um.

### Axons make intimate contacts at the tumor bed

To gain a more accurate understanding of the spatial relationship of nerve twigs and tumor cells, we double IHC stained cases of HGSOC for PAX8 (HGSOC lineage marker) [41] and β-III tubulin. In many instances, tumor-infiltrating nerves were in close proximity to PAX8 positive tumor cells (Figure 3E), suggesting intimate associations forming at the tumor bed. Recent studies demonstrate the presence of bona fide synapses in brain tumors [42-44]. While peripheral sensory nerves may not generally form synapses, they do respond to signals in the local environment by releasing factors from their nerve terminals. Importantly, among the ligands that activate TRPV1 channels are protons (low pH) which are particularly abundant in the TME suggesting that tumor-infiltrating sensory nerves may become activated by the tumor milieu [45, 46].

Whether electrical or chemical in nature, these neural connections should elicit measurable electrical activity. To test this hypothesis, patient tumor slices were electro-physiologically analyzed by microelectrode arrays (MEA). MEAs contain multiple microelectrodes that stimulate and record electrical activity from overlying cells or tissue slices [47]. Fresh tissue slices (n≥4 slices/patient sample) were generated acutely from n=7 HGSOC cases, n=5 benign gynecological tumors and n=2 normal ovaries and maintained in oxygenated artificial cerebrospinal fluid to preserve neuronal function. The MEA utilized, pMEA100/30, contains a 6×10 electrode grid where 30µm electrodes are spaced 100µm apart as depicted in Supplemental Figure 3 E, F. An image of a tissue slice within an MEA is shown in Supplemental Figure 3G. For all tissue slices, baseline activity is recorded for approximately 20 seconds, then selected electrodes are stimulated and evoked responses recorded for the next 20 seconds after which the stimulus is removed and electrical activity recorded for the final 20 seconds to assess reversion back to baseline.

While the majority of samples harbored little to no spontaneous electrical activity, stimulation of one or more electrodes induced measurable evoked responses from other electrodes. Example electrical trace recordings from malignant HGSOC (Figure 4A, B), benign gynecologic disease (Figure 4C, D) and normal ovary (Figure 4E, F) slices are shown. Here (Figure 4A, C, E), the activity of each electrode is represented by a different colored line and is recorded before (baseline), during (evoked activity) and after (reversion to baseline) stimulation. When all the recordings from malignant, benign and normal tissue slices were collected and averaged, we found no significant differences between benign and normal activity (data not shown). However, significant differences between malignant and benign tumors were noted. The mean spike amplitude before stimulation (i.e. baseline activity) was significantly higher in malignant slices as compared to benign slices (Figure 4G). Similarly, the mean spike amplitude during stimulation (i.e. evoked activity) was also significantly higher in malignant vs benign slices (Figure 4H) as was the mean amplitude difference between pre- and post-stimulation (Figure 4I). Additional slices from all samples were fixed, paraffin-embedded and histologically stained to confirm the presence of tumor (Supplemental Figure 4C-F). These electrophysiologic data show that HGSOCs are more electrically conductive than benign/normal tissue and are consistent with our hypothesis that tumor-infiltrating nerves establish functional neural circuits within HGSOCs. This elevated activity may reflect increased numbers of nerves or their level of circuit complexity in HGSOCs.

**Figure 4.**
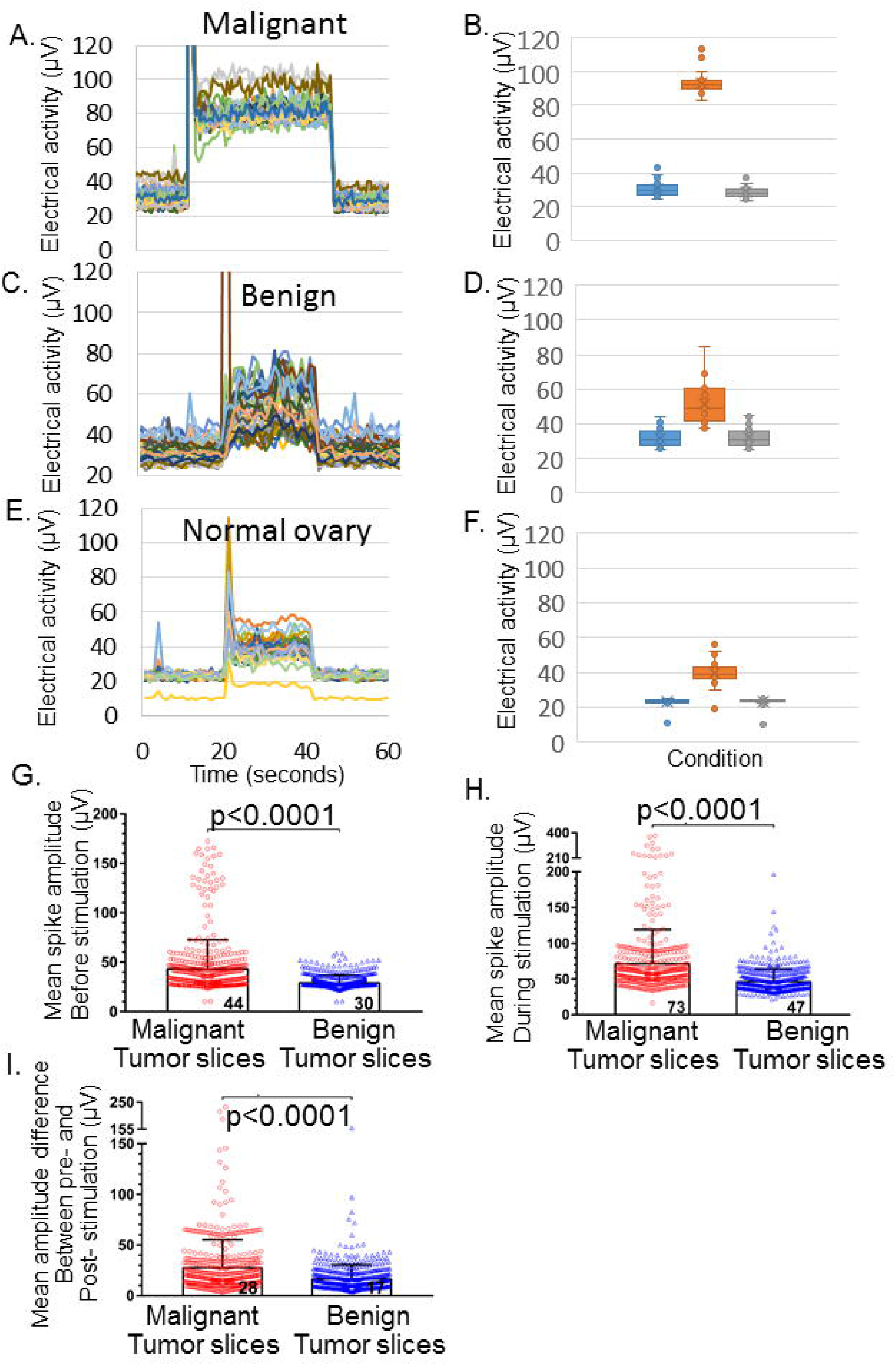
Electrical activity in acute tumor slices recorded on a MEA. MEA recordings of malignant HGSOC (A), benign (C) and normal ovary (E) slices before, during and after stimulation. Baseline activity is recorded for the first 20 seconds, followed by stimulation and recording for the next 20 seconds after which the stimulation is shut off and return to baseline is recorded for the last 20 seconds. B, D, F) Average electrical activity from all electrodes is plotted as a box and whisker plot. Blue, baseline; orange, during stimulation; grey, post-stimulation. Standard deviation, error bars. N≥4 slices/tumor were analyzed; the number of slices determined by the tumor size; n= 7 HGSOC samples, n=5 benign gynecologic tumor samples and n=2 normal ovary were analyzed. The mean spike amplitudes were calculated and only the data from electrodes with statistically significant evoked responses (p<0.01) were used in further analysis to compare slices from malignant and benign tumors as follows. G) Mean spike amplitude before stimulation. H) Mean spike amplitude during stimulation. I) Mean amplitude difference: (mean amplitude during stimulation) - (mean amplitude before stimulation). Each symbol represents an electrode: 456 electrodes for 18 slices from 7 malignant tumors and 488 electrodes for 10 slices from 4 benign tumors. Columns and bars show mean ± S.D. The numbers inside the columns are the mean values. Statistical significance was determined by the Mann-Whitney nonparametric test.

Our data indicate the presence of functional sensory neural circuits within HGSOCs. To further validate this, we tested if electrical activity could be pharmacologically blocked. Following recording of baseline and evoked activity, HGSOC slices were incubated with lidocaine, a voltage-gated sodium channel blocker, and the same slices again analyzed by MEA. Representative box and whisker plots for two different HGSOC samples show that average electrical responses before, during and after stimulation are dampened by lidocaine treatment (Figure 5 A-D). While lidocaine is predominantly considered a voltage-gated sodium channel blocker, it also functions as a TRPV1 channel sensitizer, activating the release of intracellular calcium stores [48]. This lidocaine-induced activation is followed by a desensitization phase [49], consistent with our electrophysiological findings. Taken together, these data indicate that functional circuits are present within neoplastic tissues, that malignant tumors harbor a greater extent or complexity of such connections (evidenced by enhanced electrical activity) and that this activity can be pharmacologically blocked; the effects of such blockade on disease remain to be defined.

**Figure 5.**
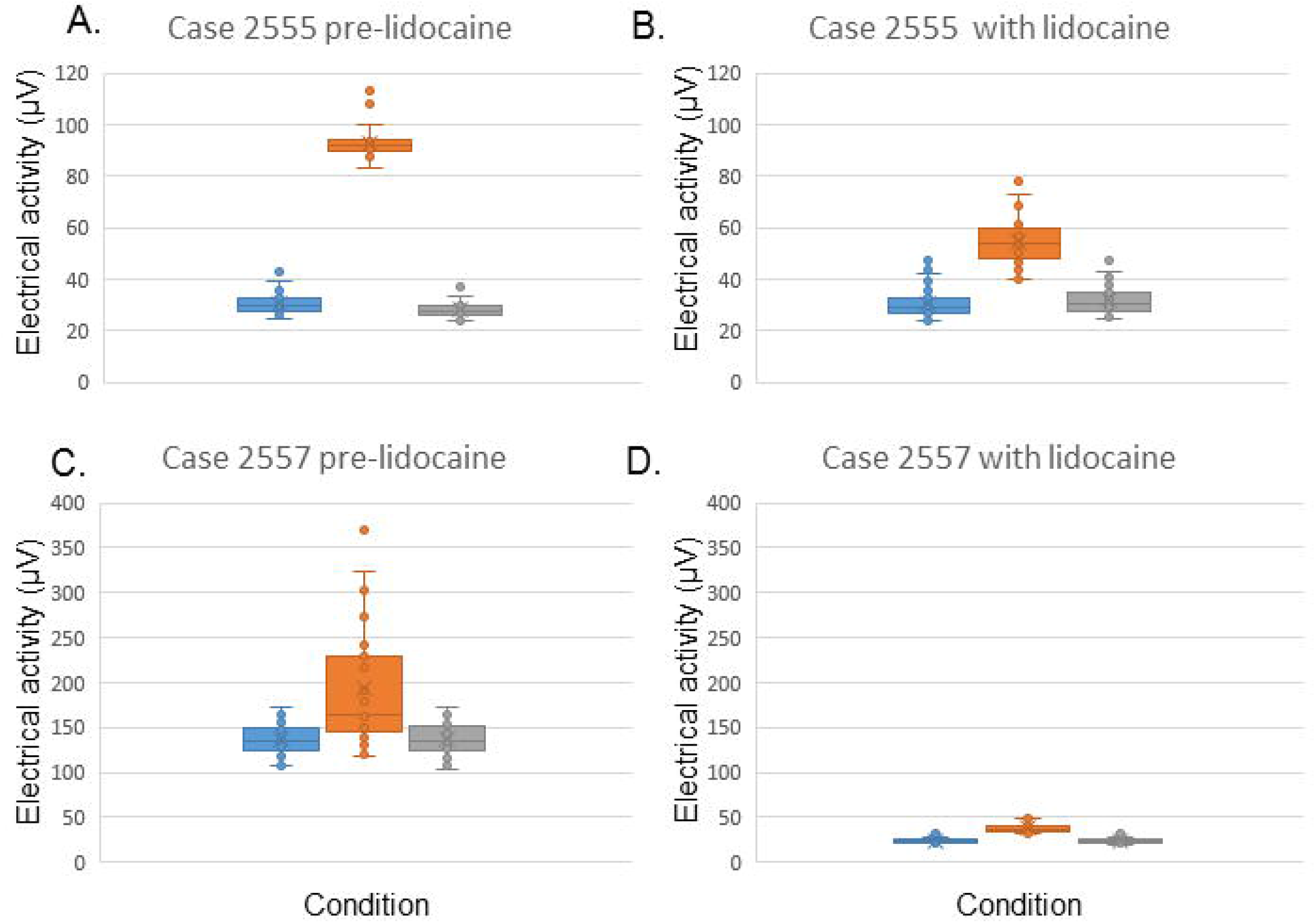
Lidocaine attenuates HGSOC electrical activity. HGSOC tumor slices analyzed as described in Figure legend 4. Box and whisker plots of average electrical responses from different HGSOC patients before (A,C) and after (B,D) lidocaine treatment (20mg/ml). Color key: pre-stimulus baseline (blue), during stimulation (orange) and post-stimulus (gray).

While these electrophysiologic data support the presence of functional neural circuits in tumor, we wondered if this activity affected survival. As an initial assessment of this possibility, we analyzed the expression of 150 neuronal-enriched genes in ovarian cancer using the OncoLnc (http://www.oncolnc.org/), Gepia2, Oncomine datasets as well as the Human Protein Atlas looking for correlations between ovarian cancer patient survival and expression of genes traditionally associated with neurons. Of the 150 neuronal-enriched genes, 47 were over-expressed in ovarian cancer; all but seven negatively correlated with survival (Figure 6A). Examples of survival plots of two such genes, Kcnt1 (a sodium-activated potassium channel) and Grid2 (a glutamate receptor), are shown in Figure 6B. When analysis specifically focused on HGSOC, a significant increase in PGP9.5 (neuronal marker) expression correlated with increasing grade (Figure 6C). Here, a series of 89 primary ovarian tumors and 36 ovarian cancer metastases (with clear pathological diagnoses) were used to establish a reference hierarchical tree. All these samples were provided by the Resource Biological Center of the Institut Curie and we processed and hybridized the chips. The dataset is publicly available on GEO (http://www.ncbi.nlm.nih.gov/geo/ under accession number GSE20565). First, the clustering was performed on this set of reference samples (89 primary tumors and 36 ovarian metastases), then the ovarian samples with ambiguous diagnosis were introduced in the dataset and the clustering was performed. These correlative patient data are consistent with a contribution of functional neural circuits to HGSOC progression.

**Figure 6.**
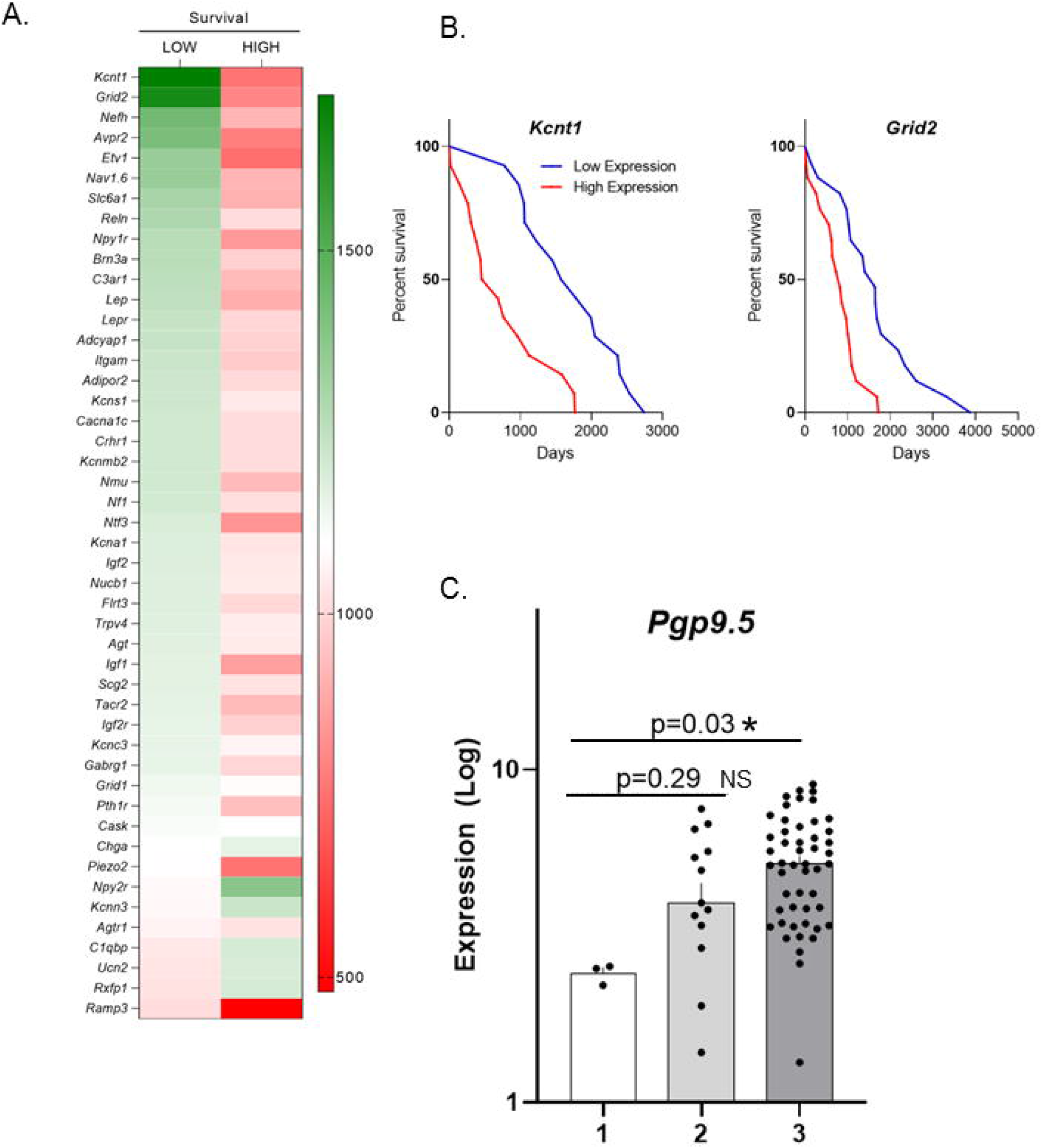
Expression of neuron-associated genes correlates with poor survival. Analysis of Oncolnc, Gepia2 and Oncomine databases for neuronal-enriched genes and ovarian cancer resulted in correlations with patient survival. A) Heatmap of neuronal gene expression and ovarian cancer patient survival. B) Kaplan-Meier survival plots of two neuronal genes (Kcnt1 and Grid2) demonstrating that elevated expression correlates with decreased survival for ovarian cancer patients. C) Clustering analysis of grade 1, 2, and 3 HGSOC samples. The clustering was first performed on reference samples (89 primary tumors and 36 ovarian metastases), then on 16 ovarian samples with ambiguous diagnosis were introduced in the dataset and the clustering was performed. n=3 for grade 1; n=15 for grade 2; n=51 for grade 3. Statistical analysis by one-way ANOVA with post-hoc Dunnet. p=0.29 (1 vs 2) ns, not significant; p=0.03 (1 vs 3), * significant.

### Tumor-infiltrating nerves contribute to tumor growth

To test of the contribution of tumor-infiltrating nerves to disease, we developed a syngeneic mouse model of HGSOC where murine oviductal secretory epithelial cells (MOSEC) from C57Bl/6 females harbor CRISPR-Cas9 mediated deletion of *Trp53* and *Pten*, commonly mutated in HGSOC [25, 38, 50, 51] (Supplemental Figure 5A). Western blot analysis of positive clones validated their retained expression of lineage markers (Pax8, Ovgp1), loss of Pten and subsequent increased expression of phosphorylated Akt (Supplemental Figure 5B). These cells generate tumors in mice that are Pax8 and WT1 positive (lineage markers) (Supplemental Figure 5C). Tumors grow following intraperitoneal (Supplemental Figure 6A) as well as subcutaneous injection (Supplemental Figure 6B) and, similar to the human disease (Figure 2A, B), these murine tumors harbor β-III tubulin/TRPV1 positive nerve twigs (Supplemental Figure 6C, D). IHC staining for neurofilament and peripherin (neuronal markers) further validate the presence of nerves in these tumors (Supplemental Figure 6E, F). Like their human counterparts, murine tumor slices respond with electrical activity upon stimulation on MEA (Supplemental Figure 7A, B). Taken together, these data support this as a faithful model of HGSOC and further suggest the presence of functional neuronal connections at the tumor bed.

To test the contribution of TRPV1 tumor-infiltrating nerves on disease *in vivo* we utilized a double transgenic mouse that lacks TRPV1 sensory nerves. This mouse is generated by crossing TRPV1-Cre and Rosa26-DTA (diphtheria toxin fragment A) mice; the resulting progeny, TRPV1-DTA, are fertile and deficient in temperature sensitivity [52]. TRPV1-DTA dorsal root ganglia lack TRPV1 immuno-positive somas confirming their genetic ablation (Supplemental Figure 7C). TRPV1-DTA or C57Bl/6 (control) female mice were subcutaneously implanted with *Trp53*^*-/-*^ *Pten*^*-/-*^ cells (1×10^5^ cells/mouse) and segregated into two cohorts. In one cohort (n=10 mice/group) we followed tumor growth and survival. Comparison of the average tumor growth curves of C57Bl/6 and TRPV1-DTA animals shows that the absence of TRPV1 nerves results in a measurable reduction in tumor growth (Figure 7A); however, this did not affect survival (Figure 7B). Nonetheless, these data suggest that tumor-infiltrating TRPV1 sensory nerves contribute to tumor growth. Individual mouse tumor growth curves can be found in Supplemental Figure 8A, B.

**Figure 7.**
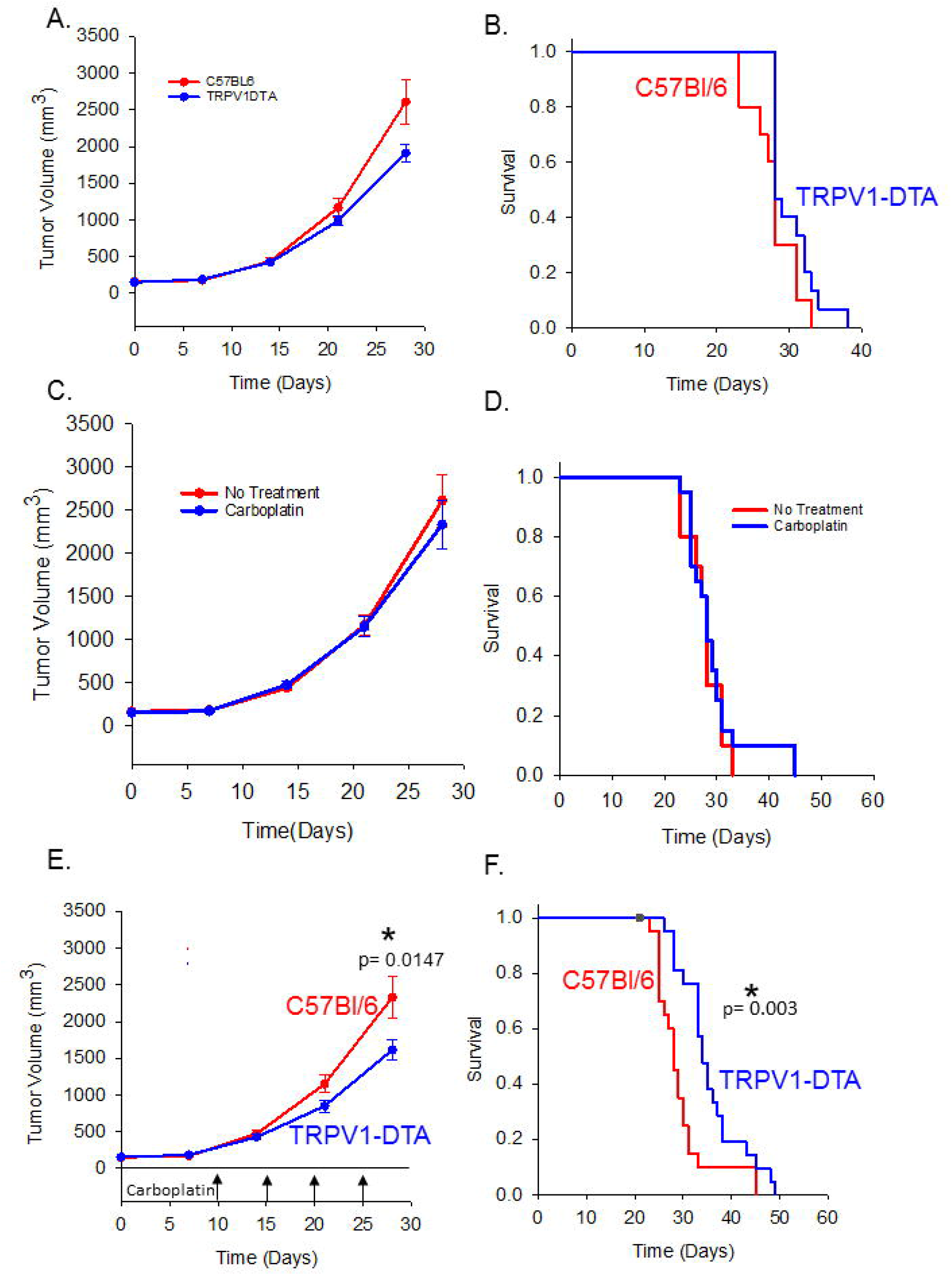
TRPV1 sensory nerves contribute to tumor growth. A) Average tumor growth curves from C57Bl/6 and TRPV1-DTA mice (n=10 mice/group) subcutaneously implanted with 1×10^5^*Trp53*^*-/-*^ *Pten*^*-/-*^ cells. Statistical analysis by one-tailed student’s t-test; *, p=0.0113, error bars, standard error of the mean. B) Kaplan-Meier survival graph of animals in panel A; statistical analysis by Log Rank test, no significant difference found. C) Average tumor growth curves of C57Bl/6 mice treated with (n=15 mice/group) or without (n=10 mice/group) carboplatin. Mice were subcutaneously implanted with 1×10^5^ *Trp53*^*-/-*^*Pten*^*-/-*^ cells. On day 10 post-tumor implantation, mice in the treatment group received weekly intraperitoneal (IP) injections with 50mg/kg carboplatin (arrows). Statistical analysis by student’s t-test, no significant difference found. D) Kaplan Meier survival plot of animals in panel C. Statistical analysis by Log Rank test; no significant difference found. E) Average tumor growth curves of C57Bl/6 and TRPV1-DTA mice (15 mice/group) following implantation with 1×10^5^ *Trp53*^*-/-*^*Pten*^*-/-*^ cells. On day ten post-tumor implantation, mice receive weekly IP injections with 50mg/kg carboplatin (arrows). Statistical analysis by one-tailed student’s t-test, p=0.0147. F) Kaplan Meier survival plot of animals in panel E. Statistical analysis by Log Rank test; *, p=0.003. Black dot, censored mouse.

### Tumor-infiltrating nerves contribute to treatment resistance

While a contribution of TRPV1 sensory nerves to ovarian cancer growth has not been previously reported, the most significant complication for HGSOC patients is treatment resistance and recurrence. In fact, treatment resistance is a nearly universal challenge for ovarian cancer patients [53]. To address this clinical issue, we focused on mice in the second cohort. Here, tumor-bearing animals (n=15 mice/group) received weekly carboplatin treatment (50mg/kg, intraperitoneal) beginning on day 10 post-tumor implantation and continuing until endpoint criteria were met. The carboplatin dose was based on a previous publication and is clinically relevant [54]. Comparison of the average tumor growth curves of treated and untreated C57Bl/6 (control) animals, demonstrates that our tumor model is resistant to carboplatin treatment as there was no effect of treatment on tumor growth or survival (Figure 7C, D). Comparison of average tumor growth curves of carboplatin-treated C57Bl/6 and TRPV1-DTA mice shows that the absence of TRPV1 sensory nerves sensitized tumor to carboplatin resulting in a significant decrease in tumor growth (Figure 7E) and a significant improvement in survival (Figure 7F). Individual mouse tumor growth curves are in Supplemental Figure 8C, D. These data suggest that tumor-infiltrating TRPV1 sensory nerves contribute to treatment resistance.

### Chemotherapy potentiates tumor innervation

These data prompted us to revisit our survey of n=10 HGSOC patient samples which demonstrated high innervation variability; we wondered if this was indicative of an underlying biology. Thus, an additional 20 HGSOC samples were collected, IHC stained, and scored. Having validated this variable innervation phenotype (Figure 1H), we wondered what clinical parameters might account for it. Standard-of-care therapy for ovarian cancer patients consists of chemotherapy; the main differences in treatment regimens involve the timing of this therapy, patients are either given chemotherapy before (neo-adjuvant) or after surgical de-bulking. We wondered whether these differences influenced tumor innervation. To assess this possibility, the previously blindly scored patient samples were separated based on naïve (no chemotherapy prior to surgery) and neo-adjuvant status. Strikingly, naïve samples (n=12) were overwhelmingly low scoring for nerve twigs while neo-adjuvant treated samples (n=18), that is residual disease, were high scoring (Figure 8A). Quantification of β-III tubulin and TRPV1 positive nerve twigs confirms the increased presence of sensory twigs in neo-adjuvant treated cases (Figure 8B) and verifies that normal ovary and fallopian tube are instead innervated predominantly by TRPV1 negative fibers. While compelling, these data were generated from unmatched samples (i.e. from different patients). Though the number of samples analyzed was relatively large (n=30), additional confirmation with matched samples was completed. Four matched cases (from the same patient) of pre-and post-treatment samples were IHC stained for β-III tubulin and scored by four independent scorers who were blinded to the sample condition. Consistent with the above finding, pre-treatment samples were low scoring for twigs while matched post-treatment samples (residual disease) were high scoring (Figure 8C). Representative photomicrographs demonstrate the striking difference in tumor-infiltrating twigs in matched samples (Figure 8F, G). These data indicate that residual disease is highly innervated and suggest that chemotherapy contributes to this phenotype.

**Figure 8.**
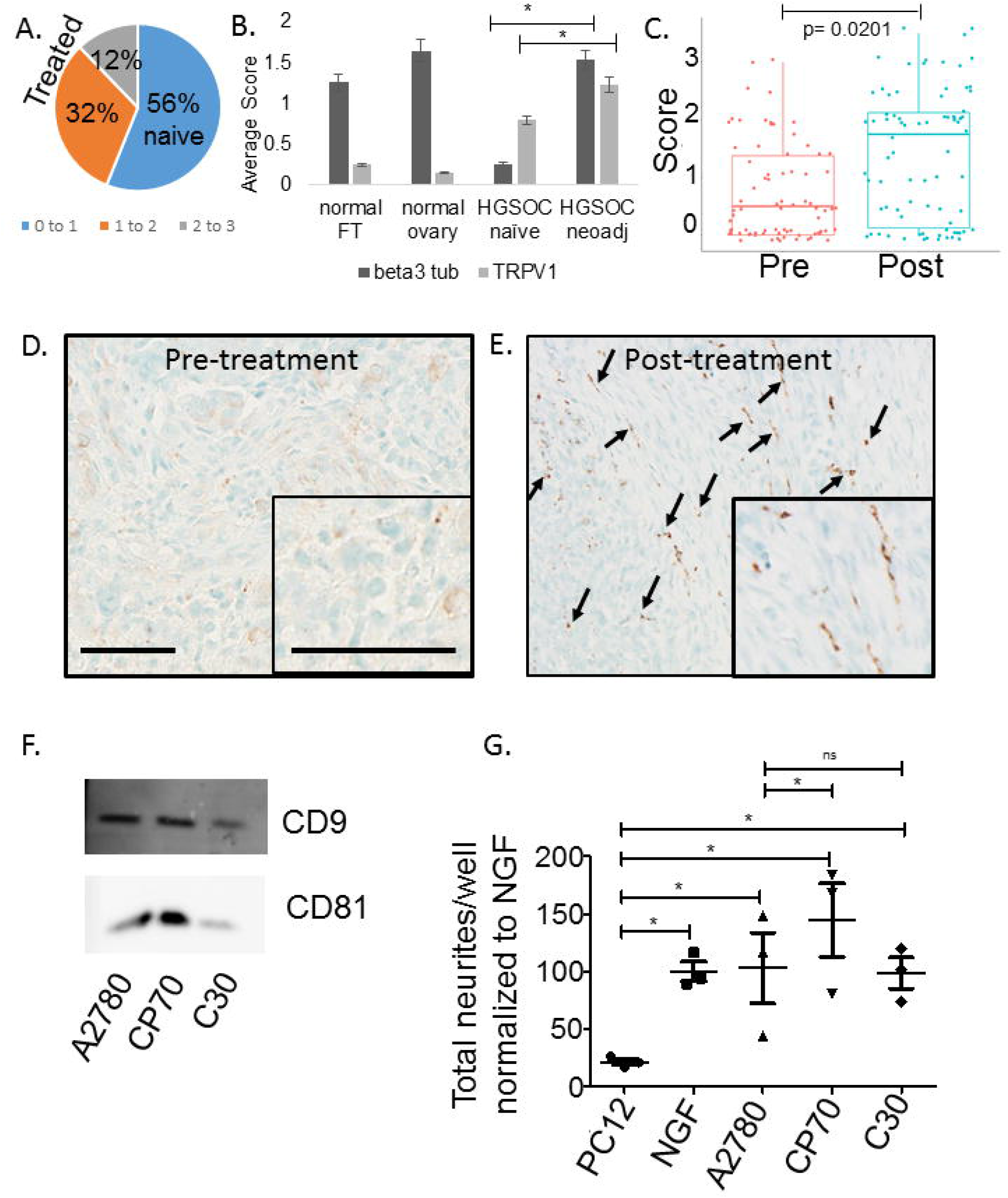
Chemotherapy increases tumor innervation. A) Pie chart of HGSOC samples showing the percent with low (0 to 1, blue), medium (1 to 2, orange) or high (2 to 3, grey) β-III tubulin IHC innervation score and the sample status: naïve (no chemotherapy before surgical de-bulking) or treated (chemotherapy treated prior to surgical de-bulking). N=30 samples were IHC stained for β-III tubulin and scored for innervation by four independent scorers, each scored n=5 random fields/sample. B) The same data from panel A showing the average twig score for β-III tubulin and TRPV1 IHC staining in unmatched HGSOC samples (n=30). Statistical analysis by student’s t-test, p<0.05. C) Innervation scores for n=4 matched (pre and post-treatment) cases of HGSOC. Scoring was as described in A. Linear mixed effects modeling was used to evaluate the change in score from pre- to post-treatment. Since the collected data consists of multiple scorers and multiple IDs, a mixed effects model was used to treat the scorer and ID as random effects. A random intercept and random slope were explored for both scorer and IDs. The random intercept allows for varying scores for each scorer and/or ID and a random slope allows for the change from pre- to post- to vary by scorer and/or ID. An indication of pre- or post-treatment score was treated as the only fixed effect. Several models were explored and compared based on differing random effects. Since each ID is rated by each scorer, the scorer factor is nested within the ID factor as a random intercept and random slope. Results of the linear mixed effects model shows a statistically significant increase in score (pre- to post-treatment) with an average of 0.8126 higher score post-treatment compared to pre-treatment score (p=0.0201). D, E) Representative photomicrographs of β-III tubulin (brown) IHC stained matched pre- (D) and post- (E) treatment tumors. Arrows, β-III tubulin nerve twigs. Insets, higher magnification. Scale bars, 50μm. F) Western blot analysis of sEVs purified from the indicated cell lines. G) PC12 cells were stimulated for 48 hours with equal amounts of sEVs purified from the indicated cells lines. Following stimulation, PC12 cells β-III tubulin immunostained and the number of neurites quantified. N=3 well/condition; experiment repeated at least n=2 times. Statistical analysis by one-way ANOVA with post-hoc Fisher’s Least Significant Difference (LSD) test was used; LSD p-values reported; * p<0.05; center value is the mean. Error bars, standard deviation. The variance between groups compared is similar.

While recurrent disease is very common with HGSOC, in many instances, it remains initially sensitive to chemotherapy. Ultimately, however, patients experience treatment resistance. Our patient data demonstrating the presence of nerves in residual disease. Our murine *in vivo* data showing a contribution of tumor-infiltrating nerves to treatment resistance. Together, these data suggest that a minimum density of tumor-infiltrating nerves is necessary to convert treatment sensitive, innervated residual disease to treatment resistant disease.

### sEVs from treatment resistant HGSOC cells have increased neurite outgrowth activity

Our data suggest that tumor-released sEVs lure nerves to the tumor bed (Figures 3D & Supplemental 3B, C) and that chemotherapy may potentiate tumor innervation (Figures 8A-G). Thus, we hypothesized that chemotherapy alters sEV cargo, endowing it with robust neurite outgrowth activity. To test this, we turned to a previously generated set of isogenic cell lines in which the parental cell line (A2780) is platinum sensitive, while two independently derived lines (C30, CP-70) are platinum resistant [55, 56]. Purified sEVs from these cell lines express EV markers (Figure 8F); equal amounts were tested on PC12 cells as previously described (Figures 3 & Supplemental 3) [18, 19]. While sEVs from the CP70 treatment resistant line induced significantly more neurite outgrowth from PC12 cells than sEVs from the treatment sensitive parental A2780 line, sEVs from the second treatment resistant line, C30, did not (Figure 8G). Interestingly, the CP70 cell line was created by intermittent exposure to increasing doses of cisplatin (a platinum drug) while the C30 line was, instead, continuously exposed to it [56]. Both *in vitro* treatment regimens produce treatment-resistant cell lines and are commonly utilized cell models to study cellular mechanisms of drug resistance [57-62]. Our data suggest that sEV cargo may be altered in different ways by these techniques.

## DISCUSSION

Here, we report on the functional contribution of tumor-infiltrating nerves to disease progression in HGSOC. Using orthogonal approaches we make four novel observations: 1) solid tumors are innervated, 2) HGSOC patient samples harbor functional neural circuits with electrophysiologic activity, 3) neo-adjuvant chemotherapy is associated with increased disease innervation and 4) tumor-infiltrating nerves contribute to treatment resistance.

Our broad assessment of innervation across common solid tumors found that all our innervated. This finding suggests that nerve recruitment to malignant disease may be a common feature in cancer. Here, we focused on HGSOC. Unlike normal ovary and fallopian tube that receive sympathetic innervation, HGSOCs are instead innervated by TRPV1 sensory nerve twigs suggesting that tumors gain nerves via active, tumor-mediated mechanisms rather than by default from native nerves in the tissue of origin. Moreover, the sensory nature of this tumor innervation suggests pain should be a key complaint from patients. The fact that the majority of ovarian cancer patients are initially asymptomatic and consequently diagnosed at late stage suggests that either nerve recruitment is a late event in disease development or that a threshold of nerves is required before pain emerges as a symptom. Interestingly, pain is one of the most prominent symptoms reported by patients that ultimately receive a diagnosis of late stage ovarian cancer [63-65]. A similar correlation between cancer pain, advanced disease and sensory innervation exists in pancreatic cancer where sonic hedgehog promotes signaling and initiation of pain via sensory nerves [66, 67]. Likewise, pain in recurrent or late stage cervical cancer remains a challenge [68-70]. As opposed to the sensory innervation we identified in cancers [18, 19], several groups have identified sympathetic and parasympathetic innervation in other tumors (prostate, liver and breast) [3, 5, 17, 71]. Recent work demonstrates varying roles for sympathetic/parasympathetic tumor-infiltrating nerves including modulation of inflammation [72], immune cell functions [17] and mediating stress effects on disease progression [73-75]. Together, these data confirm a neural contribution to cancer and emphasize the need to mechanistically define pathways promoting neural recruitment and functionally relevant intra-tumoral neural interactions that contribute to disease progression.

Towards the first end, we tested the possibility that tumor-released sEVs lure nerves to the tumor bed. We show that sEVs from ovarian cancer cell lines harbor neurite outgrowth activity and that stable expression of one oncogene (either Myc or Ras) is sufficient to endow FT33 (fallopian secretory cells) sEVs with robust neurite outgrowth capacity. We interpret these data to suggest that sEV-mediated recruitment of nerves is a critical event for disease initiation such that oncogenic transformation is sufficient to promote sEV-mediated tumor innervation. If true, tumor innervation should be an early event in cancer growth. In support of this hypothesis, Magnon *et al* show that ablation of sympathetic nerves inhibits formation of prostate cancer in a mouse model of the disease [3]. Also consistent with this hypothesis is the neural signature identified in fallopian tube precursors with loss of H2Bub1 [23]. Chemical and physical de-nervation studies by additional groups have verified the critical contributions of nerves to cancer formation [76]. Our data show that early in the process of oncogenesis, possibly upon acquisition of an oncogenic driver, sEV cargo is modified such that nerve recruitment capacity is achieved.

While defining a mechanism of nerve recruitment is important, determining how nerves contribute to disease progression would provide additional insights into the neural regulation of cancer. Support for this concept comes from our *in silico* analysis showing that expression of neuronal genes significantly correlated with higher stage disease in HGSOC. This result is bolstered by the finding that expression of neural mRNAs is unfavorable for prognosis in HGSOC [24]. We functionally tested the hypothesis that nerves form functional connections at the tumor bed by measuring electrical activity in patient samples. While we record evoked electrical activity from normal ovary, benign ovarian disease and HGSOC patient samples, the magnitude of these responses was greatest in malignant disease. These data suggest that malignant disease either potentiates nerve recruitment (as compared to normal ovary and benign tumors) or elaborates functional circuits that are more complex. Recent work by Sharma *et al* shows that during development, neurons respond to sEVs with increased proliferation (neurogenesis) and enhanced neural circuit formation [77]. Moreover, neuronal activity promotes brain growth [78]. It is not surprising then, that brain tumors utilize a similar mechanism to enhance malignant growth *in vivo* [42, 43, 79]. The usurping of developmental pathways in cancer is common; the use of sEVs to recruit nerves to the tumor bed may be a previously unappreciated reflection of sEV-regulated neurodevelopmental programs. Whether peripheral tumors similarly exploit neuronal activity to drive their growth remains to be fully tested; our MEA data demonstrating enhanced electrical activity in malignant HGSOC are consistent with such a mechanism.

An alternative interpretation of the enhanced electrical activity in HGSOC slices is that the tumor cells themselves (or other cells in the tumor microenvironment) harbor increased expression cation channels and are instead responsible for the activity we measure. In fact, epithelial cells have been found to express functional TRPV1 channels that elicit electrical activity [80, 81]. This warrants further investigation.

Importantly, the ability of lidocaine to attenuate evoked responses suggests that voltage-gated sodium channels alone or in combination with TRPV1 channels significantly contribute to the activity measured and indicate their potential use as drug targets. Our *in vivo* data demonstrating that TRPV1 nerves contribute to tumor growth as well as treatment resistance support the hypothesis that silencing these tumor-infiltrating nerves may elicit a biologically favorable response. While the possibility of quenching intra-tumoral neuronal activity as a cancer therapy has yet to be tested, our data strongly support this concept. If proven true, such a strategy may unlock the potential utility of currently FDA approved neurological drugs for cancer treatment.

It must be noted that nerves require trophic factors to remain not only functional but also functionally connected [82, 83]. While not the focus of this work, our data imply the expression of neurotrophic factors within the tumor microenvironment and suggest they represent worthy targets for therapeutic intervention that may short circuit intra-tumoral neural connections and thus, indirectly control tumor growth.

Perhaps the most intriguing discovery from this study is the contribution of chemotherapy to tumor innervation and residual disease. Using matched and unmatched patient samples, we show that neo-adjuvant chemotherapy correlates with highly innervated, residual disease. When sEVs from two independently generated platinum-resistant ovarian cancer cell lines were tested on PC12 cells, only one (CP70) demonstrated potentiation of neurite outgrowth activity, the other (C30) harbored neurite outgrowth activity similar to the platinum sensitive parental line (A2780). Understanding how the two treatment resistant cell lines were derived, sheds light on these data. The CP70 cell line was generated by intermittent exposure to increasing concentrations of cisplatin; the C30 cell line was instead produced by continuous drug exposure. Given the side effects of chemotherapies, cancer patients do not receive continuous chemotherapy; instead treatment regimens typically consist of a period of drug infusions followed by a defined “rest” (off drug) period and this pattern is repeated for a number of cycles. Thus, generation of the CP70 drug resistant cell line closely mimics clinical patient treatment protocols. While this approach is necessary, clinical trials have repeatedly demonstrated that shortening the period of time between chemotherapy infusions provides a survival advantage [84, 85]. Moreover, recurrent disease is more prevalent in patients that are neo-adjuvant treated as opposed to those that receive up-front surgical de-bulking [86]. Importantly, one study shows that neo-adjuvant therapy increases the risk of platinum-resistant recurrent disease at late stage [87]. These findings together with our data suggest that chemotherapy modulates sEV cargo such that robust innervation of residual disease ensues that may ultimately contributes to platinum-resistant recurrent disease. Changes in sEV cargo induced by chemotherapeutic agents have been previously documented and support a bystander effect of chemotherapy on sEVs that ultimately contributes to disease progression [55, 88, 89].

Taken together, our data suggest that chemotherapy modulates sEV cargo potentiating its tumor innervation capabilities, driving treatment resistance and disease progression. If correct, this hypothesis predicts that the time to treatment resistance and disease progression will be shorter in patients receiving neo-adjuvant therapy as compared to those that have primary de-bulking surgery. Published clinical trials support this prediction [87, 90]. We further validate this hypothesis with our syngeneic carboplatin-resistant ovarian cancer model; we show that simply removing TRPV1 sensory nerves (TRPV1-DTA mouse) is sufficient to sensitize tumors to carboplatin therapy and improve survival. While our findings require additional confirmation, they suggest that patients receiving neo-adjuvant chemotherapy may benefit from the addition of pharmacological agents that block exosome release and/or nerve signaling. While not currently clinically available, high-throughput screening of FDA approved drugs has already identified agents with inhibitory exosome secretion activity [91]. These drugs hold great promise for combination therapeutic approaches in oncology. Similarly, as our understanding of the neural composition of cancer expands and key neurotransmitters, channels and neurotrophic factors are identified, it is likely that FDA approved neurological drugs can be successfully repurposed for use in oncology.

## Supporting information

Kovacs et al supplementary data2

## ACKNOWLEDGEMENTS

This work was supported by the National Institutes of Health, National Institute of General Medical Sciences, Center of Biomedical Research Excellence (5P20GM103548-08), the Ovarian Cancer SPORE (1P50CA228991), the Molecular Pathology and Imaging cores at Sanford Research (supported by 5P20GM103548 and P20GM103620), the Kansas Institute for Precision Medicine COBRE (P20 GM130423), the Canadian Institutes of Health and Research (#407016) and the Ovarian Cancer Translational Center of Excellence at Penn Medicine’s Abramson Cancer Center. This work was also supported by the Honorable Tina Brozman “Tina’s Wish” Foundation, the OVERRUN Ovarian Cancer Foundation, the Dr. Miriam and Sheldon G. Adelson Medical Research Foundation, the Claneil Foundation and the Basser Cancer for BRCA. We would like to thank Ms. Ashley Tetlow for her technical assistance.

## DEDICATION

This work is dedicated to Amy Joy Cohen-Callow, PhD who faced ovarian cancer bravely and gracefully. She never gave up hope for a cure, not even when things looked quite grim. She is sorely missed yet ever present in our hearts. Amy’s unending belief that research would one day find a cure for this horrid disease fuels our continued efforts to contribute towards that end.

## AUTHOR CONTRIBUTIONS

AK: MEA, MEA analysis, review of manuscript

DWV: generation of hypothesis, designing experiments, in vivo animal studies, intellectual contributions, critical review of manuscript

MM: designing experiments, purification of exosomes, PC12 assay and quantification, scoring of IHC staining, review of manuscript

HR: MEA, MEA analysis, exosome purification, review of manuscript

SJV: scoring of IHC staining (performed with PDV), dissecting of mouse tumors for MEA, MEA analysis

CSW: procurement of human samples, review of manuscript

AR: atomic force microscopy, review of manuscript

JS: scoring of IHC staining, immunofluorescent staining of samples

CTL: DRG isolation and staining from TRPV1-DTA mice, review of manuscript

JC: CX7 analysis, review of manuscript

MB: procurement of human samples, review of manuscript

MM: procurement of human samples, review of manuscript

JYY: procurement of human samples, review of manuscript

MM: ovarian and FT cell lines, review of manuscript

NT: developed syngeneic ovarian cancer model, review of manuscript

SS: developed syngeneic ovarian cancer model, review of manuscript

AB: procurement of human samples, review of manuscript

DKO: procurement of human samples, review of manuscript

EJ: procurement of human samples, review of manuscript

LES: procurement of human samples, review of manuscript

TE: analysis of public datasets, review of manuscript

ZH: atomic force microscopy, review of manuscript

JW: CX7 analysis, review of manuscript

JEH: procurement of human samples, review of manuscript

AKG: treatment resistant cell lines, EV purification, critical review of manuscript

ST: analysis of public datasets, review of manuscript

RD: design of research studies, generation of hypothesis, analysis of data, critical review of manuscript

PDV: design of research studies, generation of hypothesis, writing of manuscript, scoring of IHC staining, MEA, in vivo animal studies, analysis of all data

## Declaration of Interests

Andrew K. Godwin is the co-founder of Sinochips Diagnostics. Ronny Drapkin is a member of the scientific advisory boards for Repare Therapeutics, Inc. and Siamab Therapeutics, Inc. and Paola D. Vermeer has a patent pending on EphrinB1 inhibitors for tumor control. Daniel Vermeer has a patent under licensing agreement with NantHealth for an HPV vaccine.

## Lead Contact and Materials Availability

Paola D. Vermeer serves as the lead contact for all materials and protocols associated with this paper. Requests for further information as well as resources and reagents may be directed to and will be fulfilled by her. All unique reagents generated by this study are also available from Drs. Vermeer and Drapkin upon completion of a Materials Transfer Agreement.

## Experimental Model and Subject Details

### Animal Studies

All *in vivo* animal studies were performed within the Animal Resource Center (ARC) at Sanford Research whose Animal Welfare Assurance is on file with the Office of Laboratory Animal Welfare. The Assurance number is A-4568-01. Sanford Health is also a licensed research facility under the authority of the United States Department of Agriculture (USDA) with USDA certificate number 46-R-009. AAALAC, Intl has also accredited the Sanford Health Animal Research Program. The ARC is a specific pathogen-free facility where mice are maintained in IVC Tecniplast Green line Seal Safe Plus cages and cages are opened only under aseptic conditions in an animal transfer station. Aseptic technique is used to change animal cages every other week. All cages have individual HEPA filtered air and animal rooms are maintained at 75^°^F, 30-70% humidity, have a minimum of 15 air changes per hour, and have a 14:10 light/dark cycle. Corncob bedding and nesting materials, both autoclaved prior to use, are maintained in all cages. Animals are fed irradiated, sterile food (Envigo) and provided acidified water (pH 2.8-3.0) *ad libitum*. There is a maximum of 5 mice/cage and they are observed daily (technicians looking for abnormal behavior, signs of illness or distress, the availability of food and water and proper husbandry). All animal experiments were performed under approved Sanford Research IACUC protocols, within institutional guidelines and comply with all relevant ethical regulations. Control (wildtype) animals injected with murine tumor cell lines were 10-15 weeks old female C57Bl/6 mice (The Jackson Laboratory), each animal was approximately 20gm in weight. Double transgenic (TRPV1-DTA) animals were generated by crossing TRPV1-Cre (The Jackson Laboratories, #017769; RRID:IMSR_JAX017769) with Rosa26-DTA mice (The Jackson Laboratory, #009669; RRID: IMSR_JAX:009669). Female progeny (TRPV1-DTA) animals were utilized at 10-15 weeks of age and were approximately 20gm in weight. n=2 C57Bl/6 and n=2 TRPV1-DTA mice were euthanized at 8 weeks of age and their dorsal root ganglia (DRG) isolated, formalin fixed and IHC stained for TRPV1 to validate absence of these nerves in double transgenic animals. Both TRPV1-Cre and Rosa26-DTA mice were backcrossed to C57Bl/6 by the depositing investigators for 10 generations and were again back-crossed by the Jackson Laboratory following deposit. Thus, the more appropriate control for the double transgenic (TRPV1-DTA) animals are wildtype C57Bl/6 mice.

All animals were randomly assigned to a cage and group. When assessing animals (e.g., measuring tumors), investigators were blinded to the groups. Animals are numbered by ear punch and cage number only. No other identifiers are on the cages to maintain investigators blinded for the duration of the experiment. When measuring tumors, investigators do not have access to the identification key.

Tumors were initiated into age-matched C57Bl/6 or TRPV1-DTA female mice as follows: using a 23-gauge needle, cells (1 × 10^5^) were implanted subcutaneously in the right hind limb of C57Bl/6 or TRPV1-DTA female mice. Caliper measurements were used to monitor tumor growth weekly. A minimum of n=10 mice/group were utilized for tumor growth studies while a minimum of n=15 mice/group were used for treatment (carboplatin) studies. Mice were euthanized when tumor volume was greater than 1.5 cm in any dimension or when other, tumor-related sacrifice criteria were met (e.g., emaciation, excessive edema, ulceration). Mice in the treatment study were treated with 50mg/kg carboplatin by intraperitoneal injection once a week starting on day 10 post-tumor implantation and continuing to the end of the experiment. When sacrifice criteria were met, mice were euthanized, tumor extracted and either fixed in neutral buffered formalin (for paraffin-embedding and IHC) or utilized fresh for micro-electrode array analysis (electrophysiological measurement).

### Human Studies

The cases for this study were obtained with patient consent and the study was approved by the Institutional Review Boards at Sanford Research, the University of Pennsylvania and Johns Hopkins. Samples from Johns Hopkins were obtained through the Legacy Gift Rapid Autopsy Program (http://pathology.jhu.edu/RapidAutopsy/). Samples from the University of Pennsylvania were obtained through Ovarian Cancer Research Center Tumor BioTrust (https://www.med.upenn.edu/OCRCBioTrust/). Ovarian cancer cases utilized in this study consisted of high-grade serous ovarian carcinoma (malignant: n=30 unmatched formalin-fixed paraffin-embedded (FFPE) tumors; n=4 matched cases; n=7 fresh tumors for MEA). Control FFPE tissues were also collected (normal ovary: n= 10; normal fallopian tube: n=10). Fresh benign gynecologic tumors (n=5) as well as normal ovary (n=2) were utilized for MEA. The benign gynecologic tumors consisted of benign mucinous and serous cystadenomas. All patients were female as males do not have fallopian tubes or ovaries and thus are not susceptible to ovarian cancer or the benign disorders mentioned above. Consented patients spanned 38-83 years of age. Formalin fixed paraffin-embedded samples were cut into 5µm sections and immunohistochemically stained.

Cases of breast, prostate, pancreatic, lung, liver and colon cancers consisted of n=10 for each cancer type. The breast cancer cases were all female and ranged in ages 43-86. The prostate cancer patient samples were all males ages 48-71. Pancreatic patient samples consisted of n= 6 females ages 48-90 and n=4 males ages 73-79. Lung cancer patient samples consisted of n=5 females ages 52-77 and n=5 males ages 54-70. Liver cancer patient samples consisted of n= 6 females ages 45-84 and n=4 males ages 56-74. Colon cancer patient samples consisted of n=5 females ages 55-85 and n=5 males ages 59-91.

### Cell lines

#### Fallopian tube cell lines

Fallopian tube secretory epithelial cells were isolated from primary human fallopian tube tissue. Fresh fimbriae were rinsed in phosphate buffered saline (PBS), finely minced and cultured for 48-72 hours at 4°C in Eagle’s Minimal Essential Medium (EMEM, Cellgro) containing 1.4mg/ml pronase (Roche Diagnostics) and 0.1mg/ml DNase (Sigma). Cultures were gently agitated during this time. Dissociated cells were incubated on Primaria plates (BD Biosciences) for 2-3 hours; this procedure removes contaminating fibroblasts and red blood cells. Non-adhered cells were seeded onto collagen-coated plates and cultured in DMEM/Ham’s F-12 1:1 (Cellgro) supplemented with 2% Ultroser G serum substitute (Pall Life Sciences) and 1% antibiotics. The purity of secretory cell culture was confirmed by immunofluorescent staining for PAX8, a mullerian lineage marker expressed by secretory, but not ciliated, cells. Additional confirmation with immunostaining for FoxJ1, a ciliated cell marker, demonstrated the absence of staining, consistent with pure secretory cell cultures. Fallopian tube secretory epithelial cells (FTSEC) were immortalized using a retroviral vector encoding the catalytic subunit of the human telomerase reverse transcriptase (*hTERT*). Increased hTERT levels were confirmed by quantitative RT-PCR. While hTERT expression prevents senescence it is unable to promote cellular proliferation and cell line expansion past approximately 10 passages. To overcome this, cells were retrovirally transduced with SV40 large T and small T antigens functionally inactivating p53 and RB1 tumor suppressor pathways. This results in enhanced growth without transforming the cells. This immortalization process generated FT33-Tag (RRID:CVCL_RK66), FT190 (RRID:CVCL_UH57), FT194 (RRID:CVCL_UH58), and FT246 (RRID:CVCL_UH61) [39, 40].

FT33-Ras and FT33-Myc cell lines were generated by transduction of FT33-Tag cells with *H-Ras*^*V12*^ or *c-Myc* respectively. Western blot analysis confirmed expression of each oncogene and retention of lineage markers [39]. Retroviral vectors used were pBABE-puro-HrasV12 and pWZL-Blast-Myc (plasmids 9051 and 10674 respectively from Addgene) and were transfected with FuGENE 6 transfection reagent (Roche Diagnostics) into HEK293T cells with medium replaced 6-12-hours later. Viral supernatants were collected 48 and 72 hours post-transfection, passed through a 0.45µm filter and applied to target cells with polybrene (8µg/ml, American Bioanalytical) for up to 24 hours. Selective antibiotics were added to the medium 72 hours post-transfection and maintained for 1 week or until cell death subsided [39].

FT33, FT190, FT194, and FT246, normal immortalized fallopian tube secretory epithelial cell lines, were cultured with DMEM:Ham’s F12 (1:1 Ratio) supplemented with 2% Ultroser G serum [39, 40]. These cell lines were authenticated using short tandem repeat profiling and tested to be free of Mycoplasma using the Cambrex MycoAlert assay at the University of Pennsylvania Perelman School of Medicine Cell Center (Philadelphia, PA) in May 2018. All FT cell lines have been deposited with ATCC.

#### Ovarian cancer cell lines

The Japanese Collection of Research Bioresources Cell Bank was the source for the following ovarian cancer cell lines: KURAMOCHI (JCRB0098; RRID: CVCL_1345) and OVSAHO (JCRB1046; RRID:CVCL_3114). ATCC was the source for the following ovarian cancer cell lines: SKOV3 (HTB-77; RRID:CVCL_0532), OVCAR3 (HTB-161; RRID:CVCL_0465), OV-90 (CRL-11732; RRID:CVCL_3768). The German Collection of Microorganism and Cell Culture GmbH was the source for the FU-OV-1 (ACC-444; RRID; CVCL_2047) cell line. The OVCAR4 (RRID:CVCL_1627) cell line was a kind gift from Dr. William Hahn’s laboratory (Dana-Farber Cancer Institute, Harvard Medical School, Boston, MA).

Kuramochi, OVSAHO, SKOV3, OVCAR3 and FU-OV-1cell lines were maintained with DMEM:Ham’s F12 (1:1 Ratio) supplemented with 10% fetal calf serum and 1% penicillin/streptomycin. OVCAR4 cells were maintained RPMI1640 supplemented with 10% fetal calf serum. OV-90 cells were maintained in 1:1 MCDB105 and Medium 199 with 10% fetal calf serum. Kuramochi, OVSAHO, OVCAR4 and FU-OV-1 cell lines are most representative of HGSOC [92].

A2780 (RRID:CVCL_0134) is a human ovarian cancer cell line derived from a patient prior to treatment (cisplatin sensitive) [93]. The CP70 and C30 cell lines are platinum resistant lines derived from the parental A2780 as follows [56]. The CP70 (RRID:CVCL_0135) cell line was generated following intermittent exposure to increasing concentrations of cisplatin (8, 20, 70 µM); the C30 (RRID:CVCL_F639) cell line was generated following continuous exposure to 30µM cisplatin. A2780, CP70 and C30 cells were maintained in RPMI1640 supplemented with 10% fetal calf serum, 100 µg/ml glutamine and 0.3 unit/ml insulin and grown at 37°C in a humidified atmosphere of 5% CO_2_ in air.

#### Ovarian cancer tumor model

The *Trp53*^*-/-*^ *Pten*^*-/-*^ murine model of HGSOC was generated as follows. *Trp53; Pten Double Knockout Murine Oviductal Secretory Epithelial Cell (MOSEC) line* The oviducts from five 6-week old C57Bl6 female mice were surgical harvested after euthanasia using a dissection microscope. Exon 5 of the *Trp53* gene and the phosphatase domain of *Pten* were targeted using the CRISPR-Cas9 system in the second passage of cultured primary MOSEC cells. The synthetic guide (sg) RNAs, GAAGTCACAGCACATGACGGAGG and TGGTCAAGATCTTCACAGAA against *Trp53* and *Pten*, respectively, were generated by annealing respective crRNA and tracrRNA pairs according to manufacturer’s instructions (Invitrogen) [94]. The cells were then transfected with the TrueCut Cas9 protein v2 (Invitrogen; Cat#A36496) and sgRNA complexes using the Lipofectamine CRISPRMAX Cas9 Transfection Reagent (Invitrogen; Cat#CMAX00008). The presence of mutations and loss of protein expression was confirmed by Sanger sequencing and Western blot analysis, respectively, in two different *Trp53*^*-/-*^; *Pten*^*-/-*^ double knockout lines (clones 2 and 4; see Supplemental Figure 5). As expected, loss of Pten resulted in phosphorylation and activation of Akt.

##### Trp53; Pten DKO tumors and tumor-derived cell lines

*Trp53; Pten* DKO MOSEC cell lines (clone 4) were expanded in culture and injected intraperitoneally (*i*.*p*.) into five 6 week old C57/Bl6 female mice (1×10^7^ cells in ice cold PBS per animal). The formation of large tumors was observed in three out of five animals 20 weeks after injections. Tumor morphology was assessed using hematoxylin and eosin staining and immunohistochemical analyses with a comprehensive panel of HGSOC markers (Pax8, OVGP1, WT-1, stathmin, pankeratin, cytokeratin 8, Ki67) and was found to be consistent with that of HGSOC. The following antibodies required a citrate buffer pressure cooker method of antigen retrieval: Pax8 antibody (ProteinTech,10336-1-AP, 1:3000), OVGP1 (Abcam, ab118590, 1:600), WT-1 (Abcam, ab89901, 1:300), Stathmin (CST, #13655, 1:150) and Ki67 (Vector Labs, VPK451, 1:1200). The following antibodies required a proteinase digestion method for antigen retrieval: PANK (DAKO, Z0622, 1:500) and CK8 (Abcam, ab154301, 1:400).

Tumor tissue isolated from tumor-bearing mice was dissociated using 90µg/ml collagenase (GIBCO, Cat#17105-041), 500µg/ml dispase (GIBCO, Cat#17105-041) and 1µg/ml DNAse I (Sigma, Cat#D4527) in culture medium (α-MEM medium supplemented with ribonucleosides, deoxynucleosides and L-glutamine (Gibco; Cat #12571-048) and containing 10ug/ml insulin-transferrin-sodium selenite (Roche; # 11074547001), 20pg/ml β-estradiol (Sigma; # E8875), 10u/ml penicillin-streptomycin solution (Invitrogen; #15140122) containing 10% fetal bovine serum (Atlanta Biologicals; Cat#S11550). Tumor-derived lines were developed and injected i.p. into ten female C57Bl6 mice. All animals developed tumors within five weeks of injection. Histological and immunohistochemical analyses of these tumors showed that they maintained HGSOC-like morphology and marker expression.

#### PC12 cells

PC12 cells were obtained from ATCC (CRL-1721; RRID: CVCL_0481) and are a rat pheochromocytoma cell line originally isolated from a male rat (*Rattus norvegicus*). PC12 cells were maintained in DMEM supplemented with 10% horse serum (Gibco) and 5% fetal calf serum (Thermofisher). When utilized in neurite outgrowth studies, PC12 cells were instead maintained in DMEM with 1% horse serum and 0.5% fetal calf serum. PC12 cells were confirmed mycoplasma free as per Uphoff and Drexler [95] at Sanford Research (Sioux Falls, SD). For all neurite outgrowth PC12 assay, n≥3 wells/condition (number of replicates depended on of the number of sEVs purified for each cell line) were utilized as technical replicates and the experiment was repeated at least n=2 times (biological replicates). Statistical analysis was by one-way ANOVA with post-hoc Fisher’s Least Significant Difference (LSD) test. LSD p-values are reported; error bars are standard deviation; center line is the mean.

### Antibodies utilized for immunohistochemistry (IHC)

Anti-β-III Tubulin (2G10, ab78078, 1:250, Abcam; RRID:AB_2256751), anti-Tyrosine Hydroxylase (Ab112, 1:750, Abcam; RRID:AB_297840), anti-TRPV1 (cat# ACC-030, 1:100, Alomone labs; RRID:AB_2313819), anti-VIP (ab22736, 1:100, Abcam; RRID:AB_447294), anti-cytokeratin (Abcam, ab8068, 1:200; RRID:AB_306238), anti-peripherin (Ab106276, 1:100, Abcam; RRID:AB_10863669), Pax8 antibody (ProteinTech,10336-1-AP, 1:3000, RRID:AB_2236705), OVGP1 (Abcam, ab118590, 1:600, RRID:AB_10898500), WT-1 (Abcam, ab89901, 1:300, RRID:AB_2043201), Stathmin (CST, #13655, 1:150, RRID:AB_2798284), Ki67 (Vector Labs, VPK451, 1:1200, RRID:AB_2314701), PANK (DAKO, Z0622, 1:500, RRID:AB_2650434) and CK8 (Abcam, ab154301, 1:400).

### Antibodies utilized for immunofluorescence (IF)

β-III tubulin Antibody (Abcam, cat# 78078, 1:100 dilution, RRID:AB_2256751), TRPV1 antibody (Alomone labs, cat# ACC-030, 1:100 dilution, RRID:AB_2313819), Synapsin1,2 (Synaptic Systems, cat#106006, 1:100 dilution, RRID:AB_2622240), PSD-95 (NOVUS, Cat# NB300-556,1:100 dilution, RRID:AB_2092366), neurofilament antibody (Biolegend, cat#837801, 1:100, RRID:AB_2565383), VGLUT1 (Synaptic Systems, cat#135303, 1:100, RRID:AB_887875).

### Antibody utilized for quantification of neurites

Anti β-III tubulin (Millipore, Ab9354, 1:1000; RRID:AB_570918). Goat anti Chicken IgY-568 (1:2000, ThermoFisher, Cat # A-11041; RRID:AB_2534098). Hoechst 33342 was used to stain nuclei (1:10000, ThermoFisher Cat# H3570).

### Nanosight Particle Tracking Analysis

The NanoSight NS300 (Malvern Panalytical, Inc., Westborough, MA) was utilized for nanoparticle (sEV) characterization. This is a laser-based system that uses light scattering and Brownian motion of particles to generate information about particle size and concentration. The NanoSight NS300 is equipped with a Blue 488nm laser and a high sensitivity scientific CMOS camera and NTA software version 3.3 (dev built 3.3.301). Each sEV sample was diluted in EV-free PBS and introduced into the NanoSight NS300 via a syringe pump that allows for a slow and constant flow of sample through the viewing chamber. Temperature is recorded and does not exceed 25°C. Approximately 30-50 particles were visualized and the camera levels were adjusted such that the particles were clearly seen but saturation was no greater than 20%. Five videos, each 60 seconds in duration, were recorded for each independent technical replicate (n=2) and all settings were maintained constant.

### Immunohistochemistry (IHC)

Tissues were obtained from the Sanford Health Department of Pathology, the BioTrust Collection (https://www.med.upenn.edu/OCRCBioTrust/) at the University of Pennsylvania and the Johns Hopkins Rapid Autopsy Program (http://pathology.jhu.edu/RapidAutopsy/). Tissues were fixed in 10% neutral buffered formalin and processed on a Leica 300 ASP tissue processor. Tissue sections were cut into 5 μm and immunohistochemically stained for β-III tubulin, TRPV1, TH and VIP; sections were also histochemically stained by hematoxylin & eosin. Antibody optimization and staining were performed with the BenchMark® XT automated slide staining system (Ventana Medical Systems, Inc.). Primary antibody was omitted as the negative control. For hematoxylin & eosin staining, slides were stained on a Sakura Tissue-Tek H&E stainer. The program runs as follows: deparaffinize and rehydrate tissue, stain in Gill’s hematoxylin (2 minutes), differentiate running tap water, blue in ammonia water, counterstain in eosin (1 minute), dehydrate and clear. For double-IHC staining, the BenchMark_®_ XT automated slide staining system (Ventana Medical Systems, Inc.) was used for deparaffinization and antigen retrieval. The antigen retrieval step was performed using the Ventana CC1 solution, which is a basic pH tris based buffer. Tissue was incubated with the antibody cocktail for 1 hour at 37 °C. Tissue was then incubated with mouse AP + rabbit HRP polymer detection kit (Biocare Mach 2 Double stain 1) for 30 minutes at room temperature. Tissues were rinsed with TBS and incubated with chromogens Betazoid DAB and Warp Red (both Biocare) for 5 minutes each, respectively. Slides were counterstained with hematoxylin, dehydrated, cleared, and coverslipped.

The Aperio VERSA 8 slide scanning system from Leica Biosystems, equipped with a Point Grey Grasshopper3 color camera for brightfield scanning was used to analyze stained sections.

### Scoring of IHC staining

Four independent evaluators (MM, JS, ET, SJV; scoring by SJV was performed in conjunction with PDV) scored all tissue samples at 20X magnification on an Olympus BX51 microscope and scored 5 random fields/sample for β-III tubulin. For HGSOC cases TRPV1 IHC staining was also scored. TH or VIP single fibers were not scored as they were scarce (unlike the presence of nerve bundles). For HGSOC scoring, the evaluators were blinded to the tissue status (naïve vs neo-adjuvant treated) while scoring. A score of 0 was given to indicate the absence of staining within each field; a score of +1 indicated 1-10% staining, +2 indicated 30-50% staining and +3 indicated greater than 50% staining. Only single nerves were scored; nerve bundles were not scored.

### Double Immunofluorescent staining

Formalin fixed and paraffin-embedded sections were deparaffinized and rehydrated by using the following washes at RT: 100% Histo-Clear (National Diagnostics) for 5min, 100% ethanol for 1 min, 90% ethanol for 1 min, 70 % ethanol for 1 min and then in PBS for 1 min. A heat-induced antigen retrieval step was performed prior immunohistochemical staining as follows: sections were incubated with 10mM Sodium Citrate Buffer (10mM Sodium Citrate Buffer, 0.05% Tween 20, pH 6.0) at 95° C for 1 hour. After cooling down at room temperature for 30min, slides were washed with PBS and then blocked in blocking buffer (1X PBS, 10% goat serum, 0.5% TX-100, 1% BSA) for 1 hour at RT. Sections were incubated with primary antibodies overnight at +4°. Slides were washed three times in PBS for 5 min each and incubated in secondary antibodies, Hoescht (1:10000, Invitrogen) at RT. Slides were washed in PBS three times, for 5 min each, and coverslips were mounted by using Faramount Mounting media (Dako). Immunostained sections were observed by using an Olympus FV1000 confocal microscope equipped with a laser scanning fluorescence and a 12 bit camera images were taken using a 60x or 100x oil PlanApo objective.

### Atomic Force Microscopy

Purified EVs were diluted 1:10 in de-ionized water and added to a clean glass dish where they were allowed to air dry for 2 hours; drying was under a gentle stream of nitrogen. EVs were then characterized using an Atomic Force Microscope (model: MFP-3D BIO™, Asylum Research, Santa Barbara, CA). AC mode was used to acquire images in air using a silicon probe (AC240TS-R3, Asylum Research) with typical resonance frequency of 70 kHz and spring constant of 2nM^-1^. Simultaneous recordings of height and amplitude images were collected at 512 × 512 pixels with a scan rate of 0.6Hz. Image processing was performed using Igor Pro 6.34 (WaveMetrics, Portland, OR) and analyzed using Image J.

### PC12 neurite outgrowth assay and β-III tubulin quantification

PC12 cells (5 × 10^4^/well) were seeded onto 96-well black optical flat bottom plates (ThermoFisher) and stimulated with 3µg of sEVs. Forty-eight hours later, cells were fixed with 4% paraformaldehyde, blocked and permeabilized with 3% goat serum, 1% BSA, and 0.5% Triton-X 100. Fixed cells were immunostained for β-III tubulin (Millipore, Ab9354; RRID:AB_570918) and nuclei stained using Hoechst 33342. Neurite outgrowth was quantified using the Cell-In-Sight CX7 High Content Analysis Platform and the Cellomics Scan Software’s (Version 6.6.0, ThermoFisher) Neuronal Profiling Bioapplication (Version 4.2). The 10x objective was used to collect twenty-five imaging fields per well with 2 × 2 binning. Hoechst positive staining identified nuclei and β-III tubulin immunolabeling identified cell somas and neurites. Cells with a Hoechst positive nucleus and β-III tubulin positive soma were classified as neurons. Analysis included only neurites longer than 20μm. All assays utilizing sEVs from cell lines were run with at least n=3 technical replicates per condition (based on sEV yield) and repeated at least two times (biological replicates) with similar results.

### sEV purification by differential ultracentrifugation

Dishes (150mm^2^) were seeded with 500,000 cells and cultured in medium containing 10% fetal calf serum which was depleted of sEVs by overnight ultracentrifugation at 110,000×g. Conditioned medium from cultured cells was harvested 48 hours later and sEVs were purified by differential ultracentifugation as described by Madeo *et al* [19]. Briefly, conditioned medium was centrifuged at 300×g for 10min at 4°C to pellet cells. Supernatants were collected and centrifuged at 2,000×g for 20min at 4°C, transferred to new tubes, and centrifuged for 30min at 10,000×g. Supernatants were centrifuged again in a SureSpin 630/17 rotor for 120 min at 110,000×g at 4°C. All pellets were washed in PBS, re-centrifuged at the 110,000xg and re-suspended in 200μL of sterile PBS/150mm dishes.

### BCA protein assay of sEVs

A modified BCA protein assay was utilized as described in Madeo *et al* [19]. Briefly, 5μl of 10% TX-100 (Thermo Scientific) were added to a 50μl aliquot of purified sEVs. This was incubated at room temperature for 10 min. A 1:11 working solution was used and incubated for 1 hour at 37°C in a 96 well plate. Absorbance (562nm) was measured (SpectraMax Plus 384) and estimates of protein concentration were generated from a standard BSA curve with a quartic model fit.

### Western blot analysis

SDS-PAGE gels (12%) loaded with equal total protein were run and transferred to PVDF membranes (Immobilon-P, Millipore). Membranes were blocked with 5% non-fat milk (Carnation) or 5% Bovine Albumin Fraction V (Millipore) and then washed with TTBS (0.05% Tween-20, 1.37M NaCl, 27mM KCl, 25mM Tris Base). Membranes were incubated in primary antibody overnight, washed and incubated with HRP-conjugated secondary antibody. Washed membranes were exposed using chemiluminescent substrate (ThermoScientific, SuperSignal West Pico) and imaged on a UVP GelDocIt 310 Imaging System equipped with a high resolution 2.0 GelCam 310 CCD camera. VisionWorksLS Image Acquisition and Analysis Software (UVP Life Science) were used to acquire and analyze images.

### Dataset Analysis

The expression of 150 neuronal-enriched genes in ovarian cancer were analyzed using Oncolnc (http://www.oncolnc.org/), Gepia2 (http://gepia2.cancer-pku.cn/#index) and Oncomine (www.oncomine.org) databases as well as the human protein atlas (https://www.proteinatlas.org/).

### Microelectrode Array (MEA)

Mouse tumors were quickly dissected and immediately sectioned using a scalpel. Fresh human tumor samples were obtained from the Sanford Health Department of Pathology or shipped overnight on ice in Miltenyi Tissue Storage Solution (cat# 130-100-008) from the University of Pennsylvania OCRC BioTrust Collection (https://www.med.upenn.edu/OCRCBioTrust/). Tumors were sectioned using a scalpel; n=4 slices were analyzed at a minimum with larger tumors allowing for a larger number of slices. n=1 slice was fixed in formalin, paraffin-embedded and stained by H&E to assess the amount of tumor present within the tissue. The approximately 900-µm-thick tumor slices were kept in oxygenated artificial cerebrospinal fluid (ACSF; 119mM NaCl, 2.5mM KCl, 1mM NaH_2_PO_4_, 26.2mM NaHCO_3_, 11mM glucose, 1.3mM MgSO_4_ and 2.5mM CaCl_2_) at room temperature. To record electrical activity, an MEA1060-Inv-BC microelectrode array system (Multichannel Systems) with a perforated microelectrode array, pMEA100/30 (Multichannel Systems), was used. pMEA100/30 has a 6×10 electrode grid and the 30-µm-diameter electrodes are spaced by 100 µm. For recordings, the tumor slice was placed on the electrodes of the pMEA and gentle suction was applied by a vacuum pump to keep the slice in place and in close contact with electrodes. Then the pMEA chamber was gently filled with oxygenated ACSF and the recording started. Electrical activity was recorded at room temperature, with 25-kHz sampling frequency, using the Butterworth 2nd order digital filter set to high pass with a cutoff frequency of 10 Hz (to eliminate slow field potentials). For electrical stimulation, a STG4000 stimulus generator (Multichannel Systems) was used. Electrical stimulation (biphasic voltage, -0.5V and + 0.5V each for 100µs and repeated after a 23ms interval) was applied to an electrode and evoked spike activities were recorded on several electrodes. Electrical activity was recorded and analyzed using the MC_Rack 4.6.2 software from Multichannel Systems.

MEA recordings from slices of: malignant HGSOC (n=7), benign gynecologic tumor (n=5) or normal ovary (n=2) were analyzed. At least n=4 slices were generated per tissue (more if sample was larger) and analyzed by MEA for a total of 25 malignant slices, 18 benign slices and 10 slices of normal ovary. Representative recordings are shown with stimulation as follows. Each plot represents recorded electrical activity over at least 60 seconds. Each tracing represents the activity from one electrode. Electrical activity is continuously recorded as follows: Baseline electrical activity is recorded for approximately 20 seconds. Selected electrodes are stimulated for a period of at least 20 seconds. The artificial stimulus is then shut off and electrical activity is recorded at least another 20 seconds during which time electrical activity reverts to baseline. Three different sets of selected electrodes were stimulated during three consecutive rounds of recordings. The first set of stimulated electrodes consisted of electrodes on the outer edges in a checker board pattern (n= 14 total electrodes stimulated); the second set of stimulated electrodes were all electrodes on the top and bottom rows (n= 20 total electrodes stimulated) while the third set of stimulated electrodes consisted of the columns of electrodes on the outer edges (n= 12 total electrodes stimulated). Some electrodes were grounded due to excessive noise. Box and whisker plots were generated and reflect the average electrical activity before, during and after stimulation for each slice. Lidocaine treatment consisted of incubation in 20mg/ml oxygenated lidocaine (Hospira, NDC 0409-4277-17) at room temperature.

### Dorsal Root Ganglia (DRG) Isolation

N=2 C57Bl/6 and n=2 TRPV1-DTA 8 week old, approximately 18gm mice were euthanized by CO_2_ and cervical dislocation. The fur was sprayed with 70% ethanol and dorsal fur removed to expose the spinal column. Standard scissors were used to remove tissue and cut the ribs leaving only the spinal column intact (head and tail were cut and removed). Laying the ventral side of the spinal column face up, incisions were made along the left and right sides all along the length of the column; once completed, the ventral half of the spine was lifted off, exposing the spinal marrow which was also removed. This exposed the DRG which were carefully removed from the surrounding tissue and placed into formalin. Following fixation, DRG were paraffin-embedded, cut and IHC stained as described.

### Statistics

Graphpad Prism V7 was used to graph and analyze PC12 neurite outgrowth data. One-way ANOVA with post-hoc Fisher’s Least Significant Difference test were utilized for statistical analysis as indicated in the figure legends. Sigma Plot (version 13) was used for graphing murine *in vivo* tumor growth and Kaplan-Meier Survival plots; Log rank test was utilized for survival analysis while student’s t-test was used for tumor growth analysis with standard error of the mean as error bars. For statistical analysis of matched pre- and post-treatment samples, linear mixed effects modeling was used to evaluate the change in score from pre- to post-treatment. Since the collected data consists of multiple scorers and multiple IDs, a mixed effects model was used to treat the scorer and ID as random effects. A random intercept and random slope were explored for both scorer and IDs. The random intercept allows for varying scores for each scorer and/or ID and a random slope allows for the change from pre- to post- to vary by scorer and/or ID. An indication of pre- or post-treatment score was treated as the only fixed effect. Several models were explored and compared based on differing random effects. Since each ID is rated by each scorer, the scorer factor is nested within the ID factor as a random intercept and random slope.

## Notes

### Summary of Updates

Patients with densely innervated tumors do poorly as compared to those with sparsely innervated disease. Why some tumors heavily recruit nerves while others do not, remains unknown as does the functional contribution of tumor-infiltrating nerves to cancer. Moreover, while patients receive chemotherapeutic treatment, whether these drugs affect nerve recruitment has not been tested. Using a murine model of ovarian cancer, we show that tumor-infiltrating sensory nerves potentiate tumor growth, decrease survival, and contribute to treatment resistance. Furthermore, matched patient samples show significantly increased tumor innervation following chemotherapy. In vitro analysis of tumor-released extracellular vesicles (sEVs) shows they harbor neurite outgrowth activity. These data suggest that chemotherapy may alter sEV cargo, endowing it with robust nerve recruiting capacity.

