## Supplementary material for "Tumor-infiltrating nerves create an electro-physiologically active microenvironment and contribute to treatment resistance": Kovacs et al supplementary data2

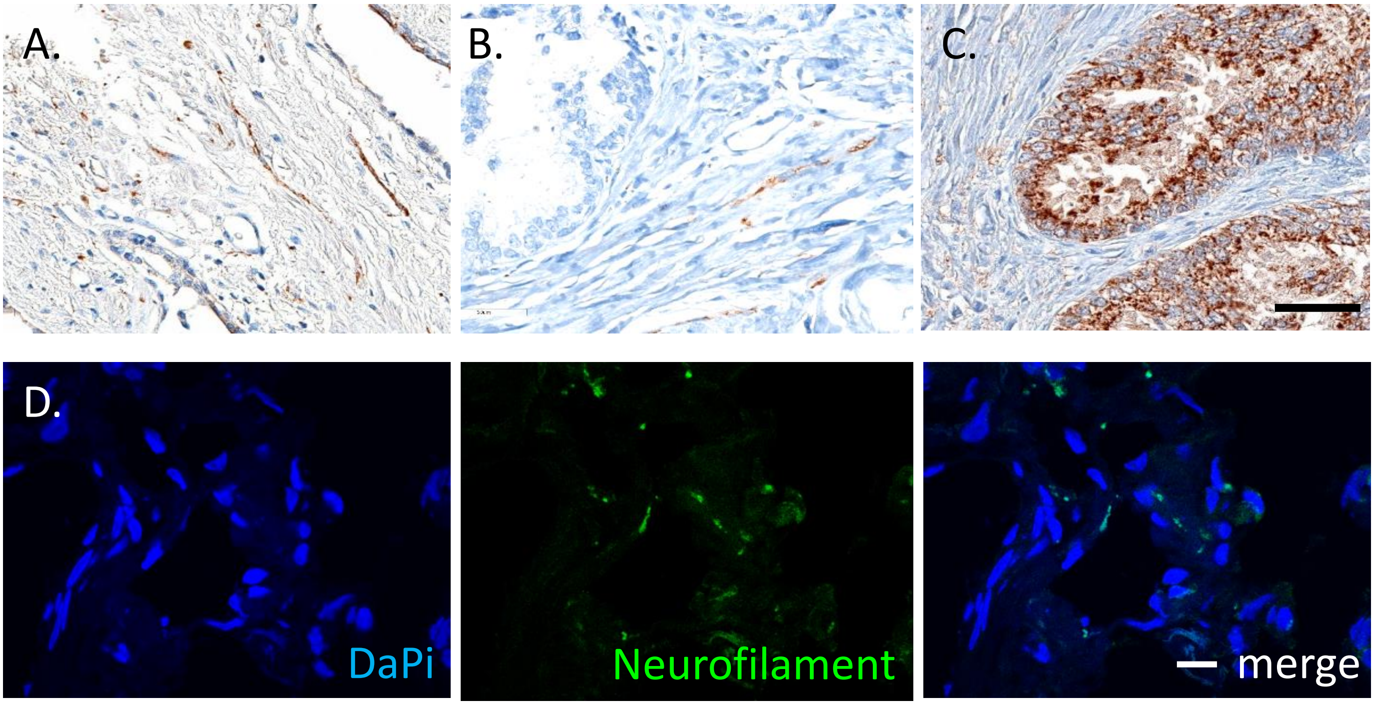


**Supplemental Figure 1.** Positive tissues for IHC staining as follows. Human prostate cancer as positive control for (A) VIP, (B) tyrosine hydroxylase, and (C) TRPV1. Scale bar, 50 µm. D) Human HGSOC immunofluorescently stained for neurofilament (green); counterstained with DaPi (blue). Scale bar, 10µm. Brightness was increased on all images in all lasers; these changes were made to the entire image.

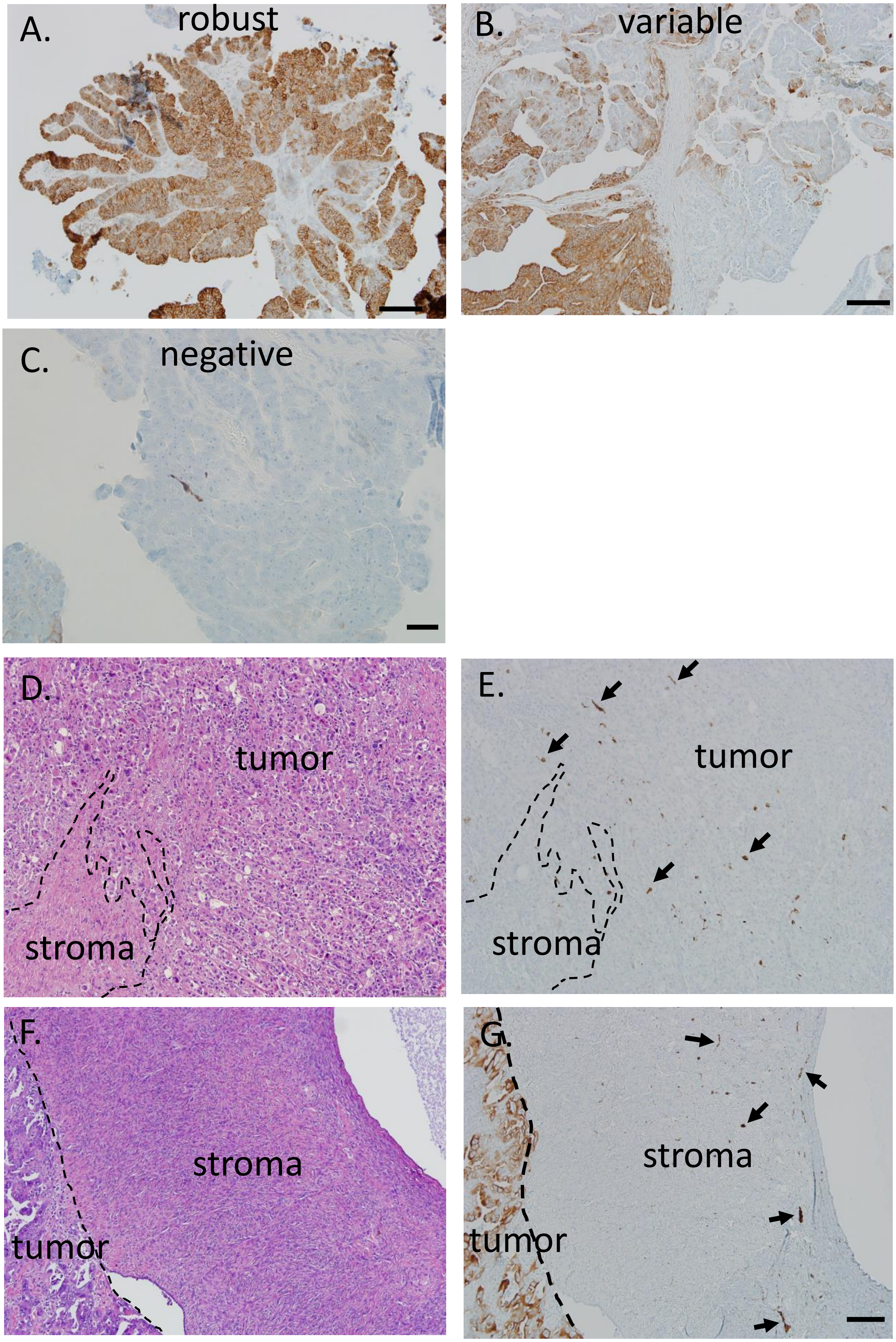


**Supplemental Figure 2.** A-C) Bright field images representative of robust, variable and negative IHC staining for β-III tubulin in tumor cells. D, E) Representative bright field images of serially sectioned HGSOC showing H&E (D) and β-III tubulin IHC staining (E, brown); dotted line outlines stroma/tumor boundary. In this example, β-III tubulin positive nerve twigs (arrows) are infiltrating directly into tumor. F,G) Representative images of serial sections from HGSOC patient stained by H&E (F) and β-III tubulin IHC (G, brown). Dotted line shows tumor/stroma boundary. In this example, β-III tubulin positive nerve twigs (arrows) are found within the stroma, in close proximity to tumor. Scale bar, (A,B,D,E,F,G) 100 μm; Scale bar, (C) 10 μm. Arrows, β-III tubulin positively stained nerve twigs.

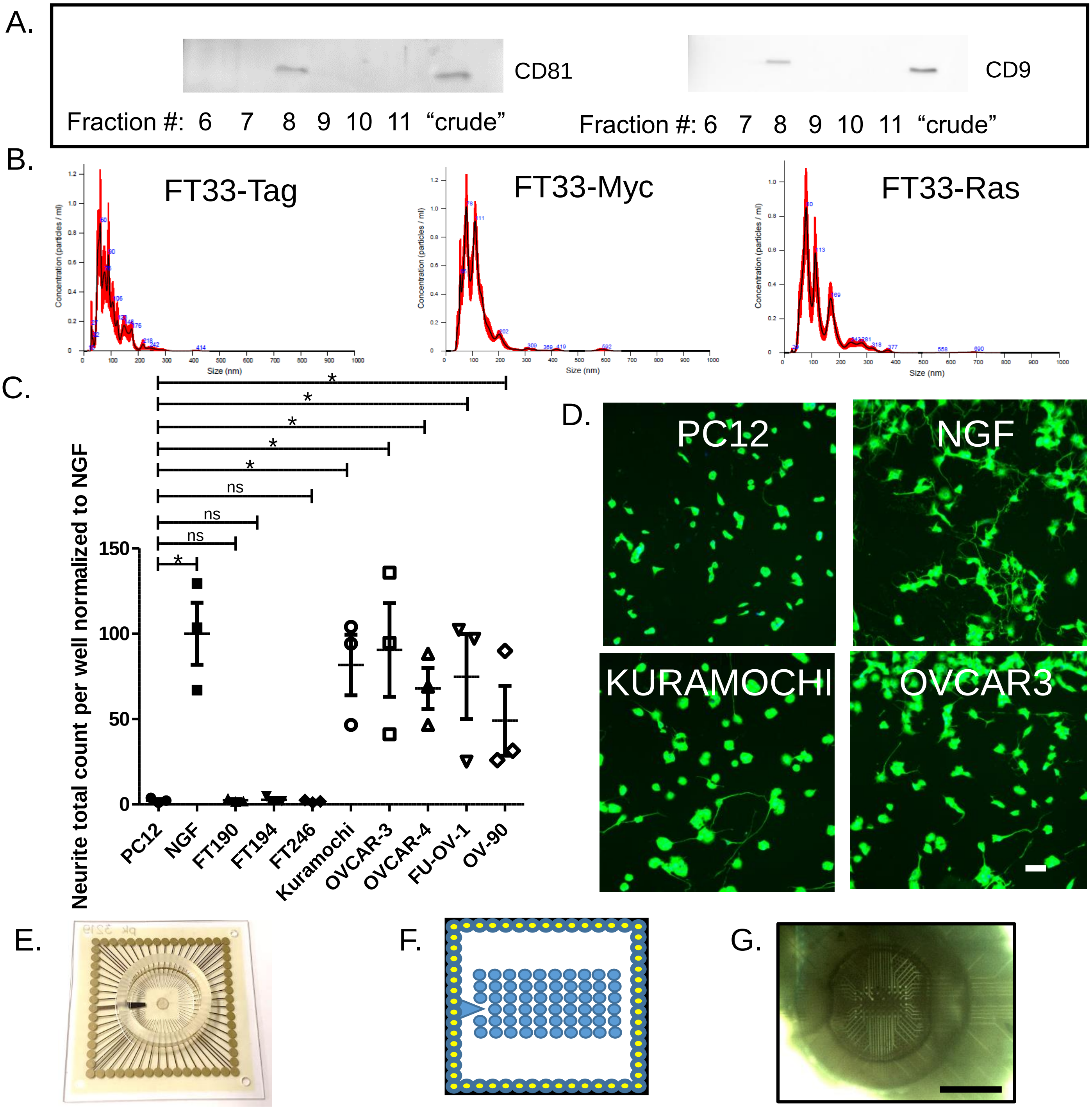


**Supplemental Figure 3.** Western blot analysis for CD9 and CD81 of optiprep density gradient fractions (fractions 6-11) of sEVs from the Kuramochi ovarian cancer cell line. "Crude" sample is that of sEVs purified by differential ultracentrifugation alone. B) Nanoparticle tracking analysis of sEVs purified from the conditioned media of FT33-Tag, FT33-Myc and FT33-Ras cells. C) Quantified neurite outgrowth activity of PC12 cells following stimulation with sEVs from the indicated cell lines. One way ANOVA with post-hoc Fisher's Least Significant Difference (LSD) test was used for statistical analysis. LSD p values reported. Center value reported is the mean. Error bars indicate standard deviation. *, $p<0.05$; ns, not significant. N=3 technical replicates/condition; experiment repeated at least n=2 (biological replicates). The variance between groups compared is similar. D) β-III tubulin immunofluorescent images of PC12 cells following stimulation with sEVs purified from the indicated cell lines. NGF treatment serves as the positive control while untreated PC12 cells (PC12) serve as the negative control. Scale bar, 40μm. E) Photomicrograph of a microelectrode array (MEA). F) Schematic representation of an MEA showing the 6x10 electrode grid (blue circles) in the center of the array. G) Photomicrograph of tumor slice on an MEA. Scale bar, 1000μm.

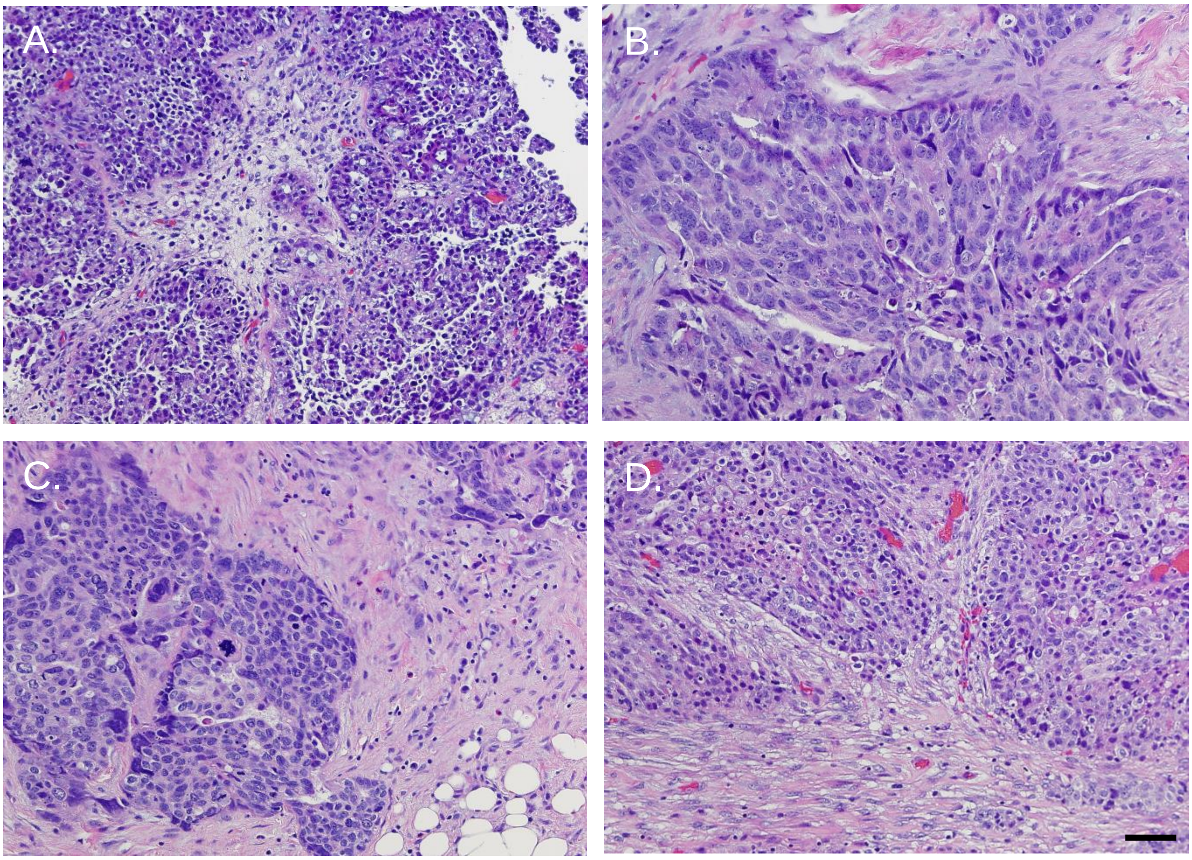


**Supplemental Figure 4.** A-D) Representative bright field images of hematoxylin and eosin stained HGSOC patient samples used for MEA analysis. All slices confirm the presence of cancer cells. Scale bar, 50µm.

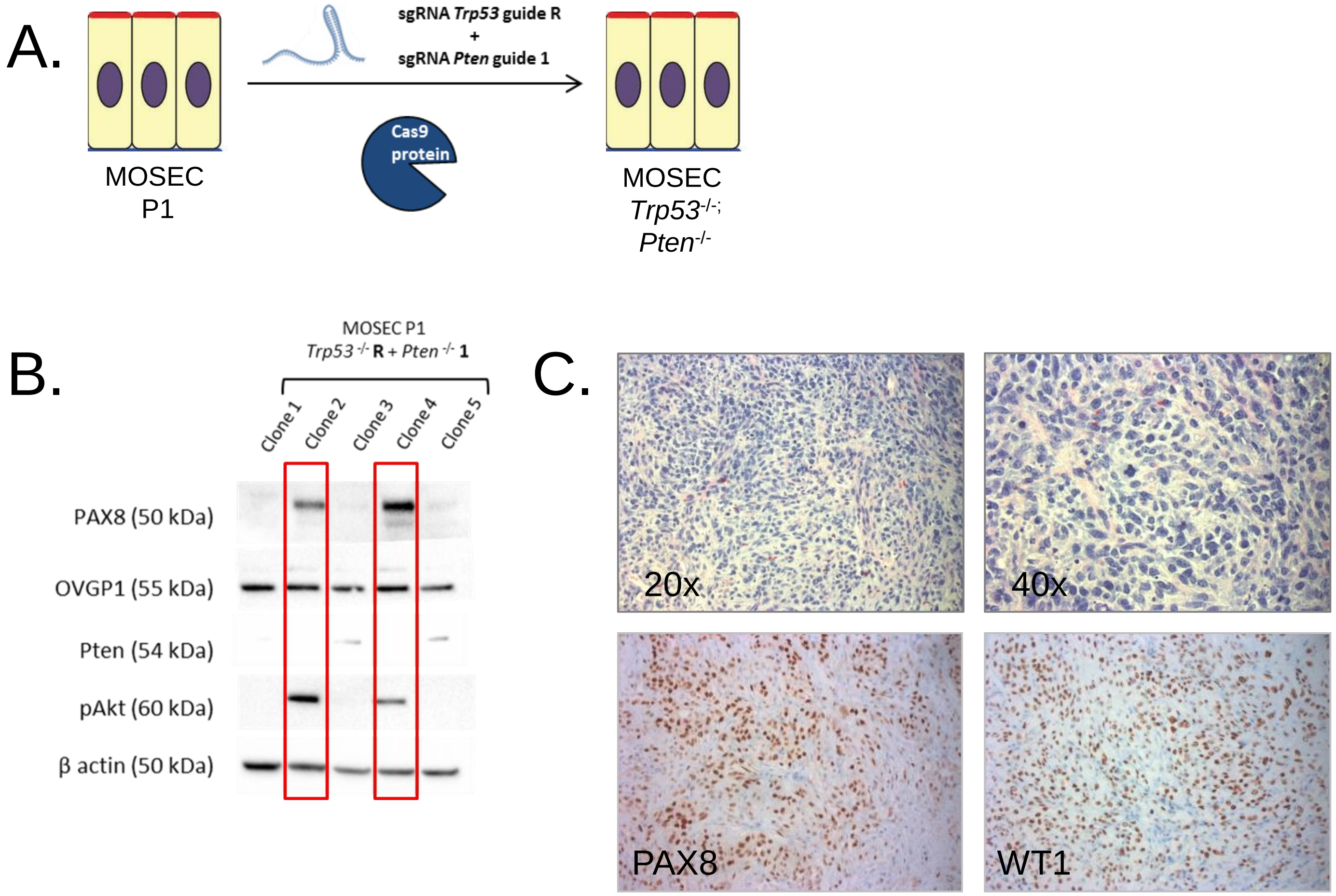


**Supplemental Figure 5**. A) Graphic illustration depicting methodology used to generate a MOSEC line deleted for *Trp53* and *Pten*. CRISPR-Cas9 was used to delete *Trp53* and *Pten* in early passage primary MOSECs. B) Western blot analysis of CRISPR-Cas9 mediated knockout of Trp53 and Pten. Five clones were evaluated for protein expression, including Pax8, Ovgp1, Pten, phospho-AKT, and β-actin. Loss of Pten protein was associated with acquisition of phospho-AKT in clones 2 and 4. C) Morphology and immunophenotype of Trp53; Pten double knockout tumors. Top panel: H&E of tumors (20x and 40x) show a morphology consistent with a high-grade carcinoma. Lower panel: immunohistochemistry for Pax8 and WT1 (20x) show that tumors retain lineage markers associated with high-grade serous carcinomas.

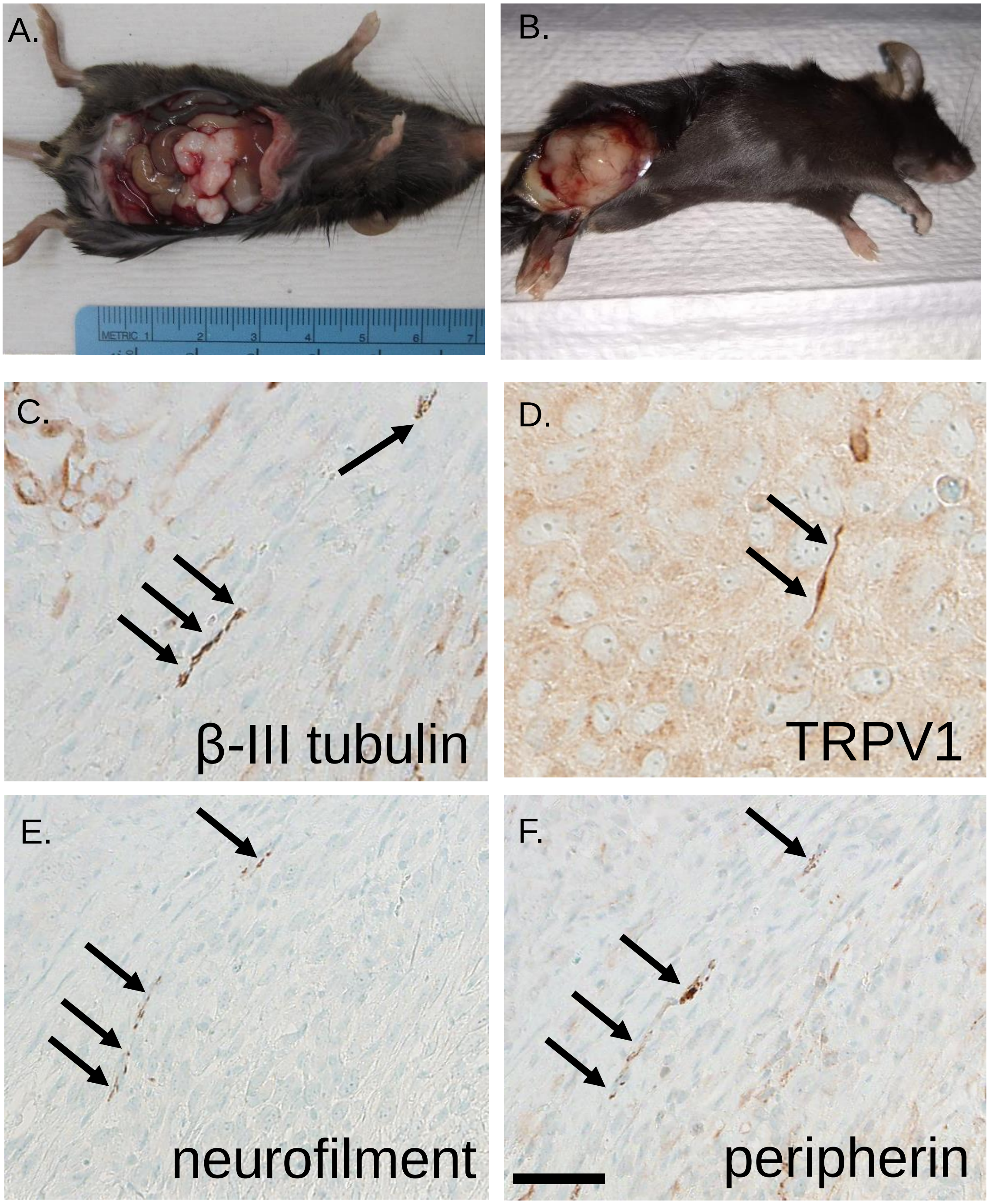


**Supplemental Figure 6.** A) *Trp53*$^{-/-}$ *Pten*$^{-/-}$ syngeneic ovarian tumors grow in the peritoneal cavity (A) as well as subcutaneously (B). Immunohistochemical staining of *Trp53*$^{-/-}$ *Pten*$^{-/-}$ tumors harbor β-III tubulin (C), TRPV1 (D), neurofilament (E), peripherin (F) positive nerve twigs (arrows). Scale bar, 50μm.

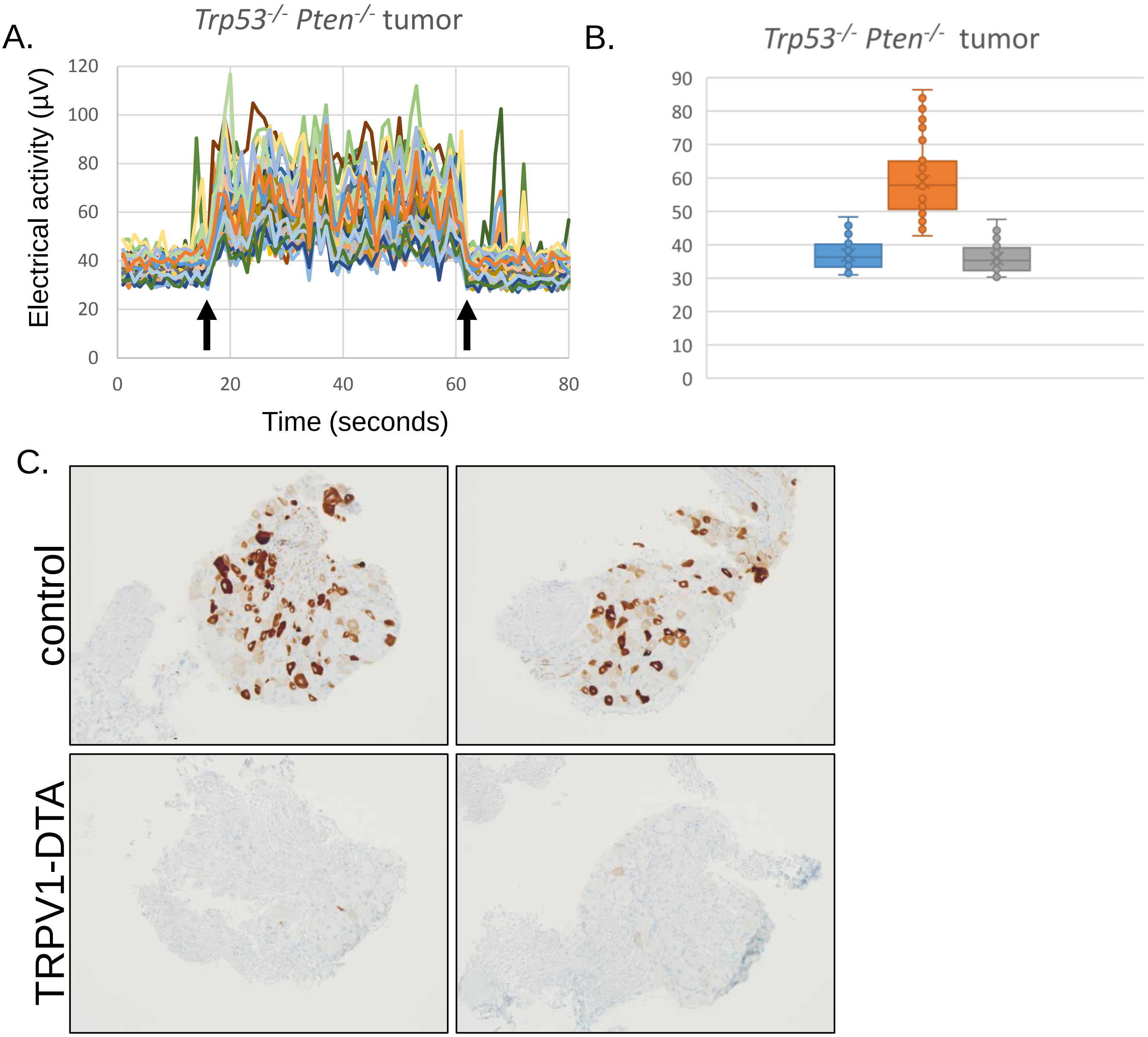


**Supplemental Figure 7.** A) Representative electrical stimulation-induced spike activity in a syngeneic *Trp53*$^{-/-}$ *Pten*$^{-/-}$ tumor slice recorded on a microelectrode array (n=12 slices, n=3 tumors). B) Box and whisker plots of the average activity before (blue), during (orange) and after (gray) stimulation. C) Representative bright field photomicrographs of dorsal root ganglia isolated from C57Bl/6 female mice (control, n=2) or TRPV1-DTA female mice (n=2) immunohistochemically stained for TRPV1 (brown).

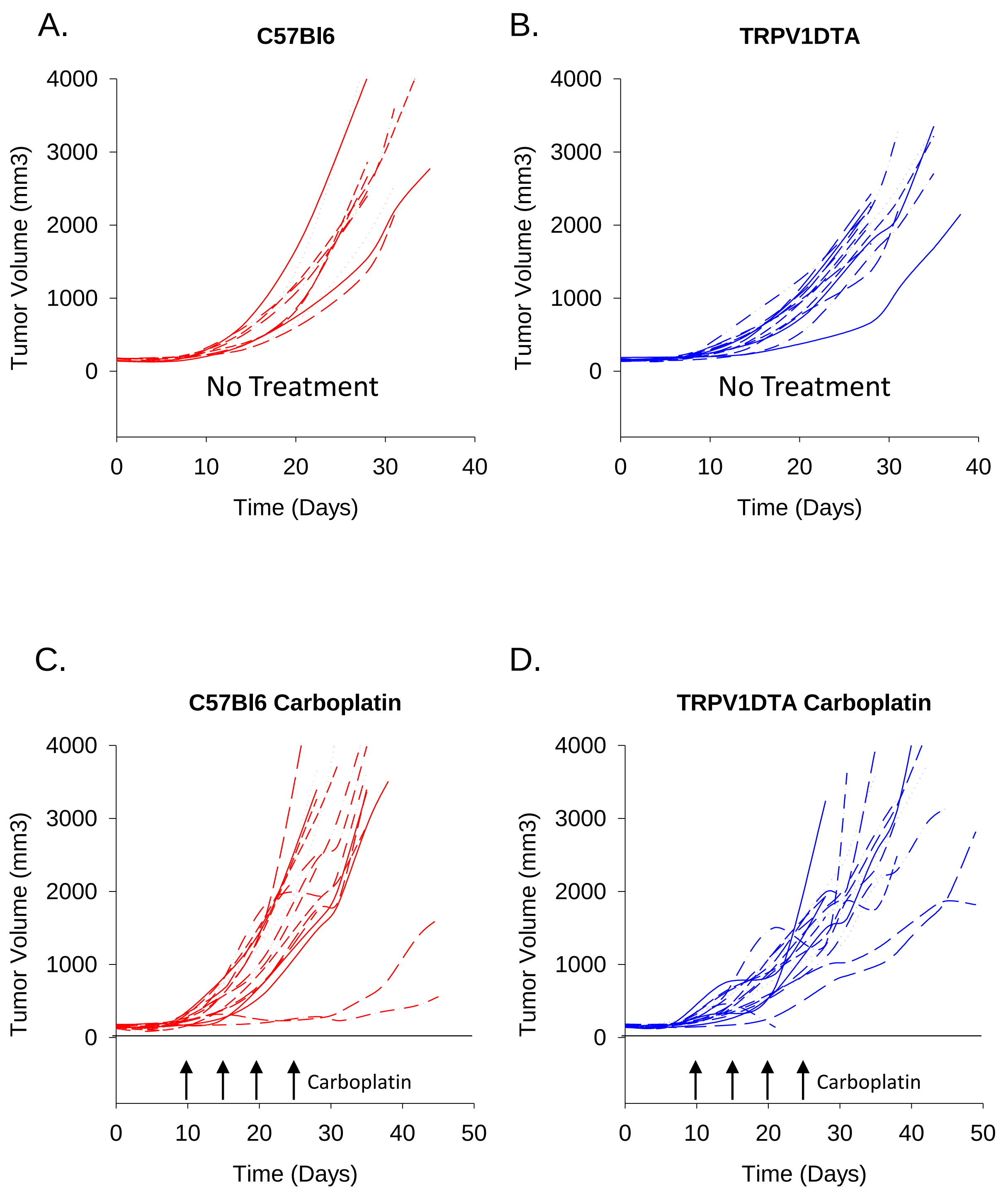


**Supplemental Figure 8**. Tumor growth curves for age-matched individual female mice bearing *Trp53*$^{-/-}$ *Pten*$^{-/-}$ tumors. A) C57Bl/6 untreated mice (n=10), B) TRPV1-DTA untreated mice (n=10), C) C57Bl/6 carboplatin-treated mice (n=15) and D) TRPV1-DTA carboplatin treated mice (n=15). Carboplatin treatment consisted of intraperitoneal administration of 50mg/kg carboplatin once a week (arrows) starting on day 10 post-tumor implantation.
